# Ancient genomes reveal Avar-Hungarian transformations in the 9th-10th centuries CE Carpathian Basin

**DOI:** 10.1101/2024.05.29.596386

**Authors:** Dániel Gerber, Veronika Csáky, Bea Szeifert, Noémi Borbély, Kristóf Jakab, György Mező, Zsolt Petkes, Frigyes Szücsi, Sándor Évinger, Csilla Líbor, Piroska Rácz, Krisztián Kiss, Balázs Gusztáv Mende, Béla Miklós Szőke, Anna Szécsényi-Nagy

## Abstract

During the Early Medieval period, the Carpathian Basin witnessed significant demographic shifts, notably under the Avar dominance for approximately 250 years, followed by the settlement of early Hungarians in the region during the late 9th century CE. This study presents the genetic analysis of 296 ancient samples, including 103 shotgun-sequenced genomes, from present-day Western Hungary. By employing identity-by-descent (IBD) segment sharing networks, this research offers detailed insights into the population structure and dynamics of the region from the 5th to the 11th centuries CE, with specific focus on certain microregions. Our evaluations reveal spatially different histories in Transdanubia even between communities in close geographical proximity, highlighting the importance of dense sampling and analyses. Our findings highlight extensive homogenisation and reorganization processes, discontinuities between Hun, Avar, and Conquest period immigrant groups, alongside the spread and integration of ancestry related to the Hungarian conquerors.

**Teaser:** Genetic formation of medieval Hungary, discontinuity between immigrant groups in the first millennium CE

## Introduction

The population history of the Migration period in Central Europe was marked by broad-scale complexity, influenced by various Eastern and Western cultural and biological impacts, as documented in written sources and archaeological studies[1], [2], [3], [4], [5], [6], [7]. In recent years, the Carpathian Basin (including the territory of today’s Hungary) has been the target for many archaeogenomic studies[8], [9], [10], [11] shedding step-by-step light on population events of this period.

Between the fall of the Roman Empire in Pannonia (433 CE) and the arrival of the Avars in 568 CE, the Carpathian Basin was home to transient groups and polities, such as the Suebs, Goths, Lombards, and Gepids, with diverse ethnic and cultural backgrounds. During this period, the Avar elite of Northeast-Eurasian origin[9], [10], [12], along with various East-European groups, settled in the region[4]. According to the written and archaeological sources, the population of the Avar Khaganate was extremely culturally diverse[5], [13]. A 20-year conflict with the rapidly expanding Carolingian Empire disintegrated the Avar Khaganate by 811 CE, which fell into smaller regional units. The former Pannonian provinces, *i*.*e*. Transdanubia (corresponding to the territory of today’s Western Hungary) and the Drava-Sava Interfluve, became a part of the Carolingian Empire, while the Danube-Tisza Interfluve and the Trans-Tisza regions (the latter two hereafter referred to as the Great Hungarian Plain, abbreviated as GHP) remained under the supervision of the Khagan. The Khaganate’s former territories had been occupied by the Moravians in the northwest and by the Croatians in the southwest, both under the supervision of the Carolingian emperor. This expansion of the Carolingian Empire was followed by the spread of its cultural influence and strategic integration of the corresponding regions during the 9th century CE. This involved the establishment of smaller, isolated counties as administrative units and a deliberate effort to Christianize the people of the newly acquired territories[5].

The power center of Transdanubia was founded at Zalavár-Vársziget (hereafter Zalavár) by Priwina and Chezil around 840 CE. Its population was presumably supplemented by communities arriving from various regions from the Baltic to the Black Sea. Between 840 and 870, three churches were built here, where the elites and their servants were being buried. Around 880 CE, the settlement became a royal center with a culturally diverse population. Around 896 CE, Prince Braslav was charged by King Arnulf with reinforcing the settlement due to the arrival of the Hungarian conquerors (also referred to as Early Mediaeval Magyars). These new arrivals overran Transdanubia in the 10th century CE and settled across the region, presumably displacing only the ruling class of the former population. Zalavár remained a prominent settlement and was recognized as the capital of both Somogy and Zala counties, which still exist today. The population density increased, and a Benedictine church was consecrated in 1019 CE, with a cemetery around it used in the following decades (further information in Supplementary Information section 1).

The Carpathian Basin faced a major cultural transition when the Hungarian conquerors arrived in the region and formed the Hungarian Kingdom by the end of the 10th century CE. The entire 10th century is referred to as the Conquest period. During this time, all previous administrative structures were disregarded, and new centers were established[5]. The extent to which the 9th century CE population survived, and either coexisted or admixed with the new conquerors, remains uncertain. These conquerors, along with the local population, were ultimately Christianized during the Árpádian era (11th-13th centuries CE).

### Research aims and sampling foci

The majority of the ancient DNA samples published from the 6th to 11th centuries CE originate from the GHP[9], [10], whereas Transdanubia almost completely lacks ancient genomes from this period, despite its significant historical events after the 5th century CE. Our primary objective was to achieve a more comprehensive genetic dataset of the area’s population, with a particular focus on the 8th to 11th centuries CE. This was undertaken to test for population continuity, as well as to establish the general structure and connections of the prevailing communities during the transition from the late Avar Khaganate to the beginning of the Árpádian era. We aimed to analyze the diversity and stability of the local population across the Carpathian Basin and the dynamics of admixture from Avar and Hungarian conquest event-related immigrants.

Overall, 294 samples were collected from various sites in Transdanubia (see Fig. 1.a, Table 1 and Supplementary Information section S1, Table S1), with special attention given to Zalavár, the Székesfehérvár area, and Visegrád (Visegrád-Sibrik hill and Visegrád-Széchenyi street 25 sites), as each represent key aspects of the studied historical transitions. The Zalavár site, known as the greatest center under the name *Mosaburg* during the Carolingian era, was sampled across three consecutive archaeological horizons: the 9th century (ZVI), the 10th century (ZVII), and the 11th century (ZVIII). Székesfehérvár, a newly formed settlement in the 10th century CE, became the royal center of the Hungarian Kingdom. The population context for the preceding centuries is represented through sampling of neighboring sites (Supplementary Information section 1). Himod-Káposztás (hereafter Himod) represents a transitional smaller rural settlement, which was sampled from two horizons (9th and 10th-11th centuries CE), while Visegrád was a newly formed religious and political center of the Kingdom in the 11th century CE.

**Table 1.**
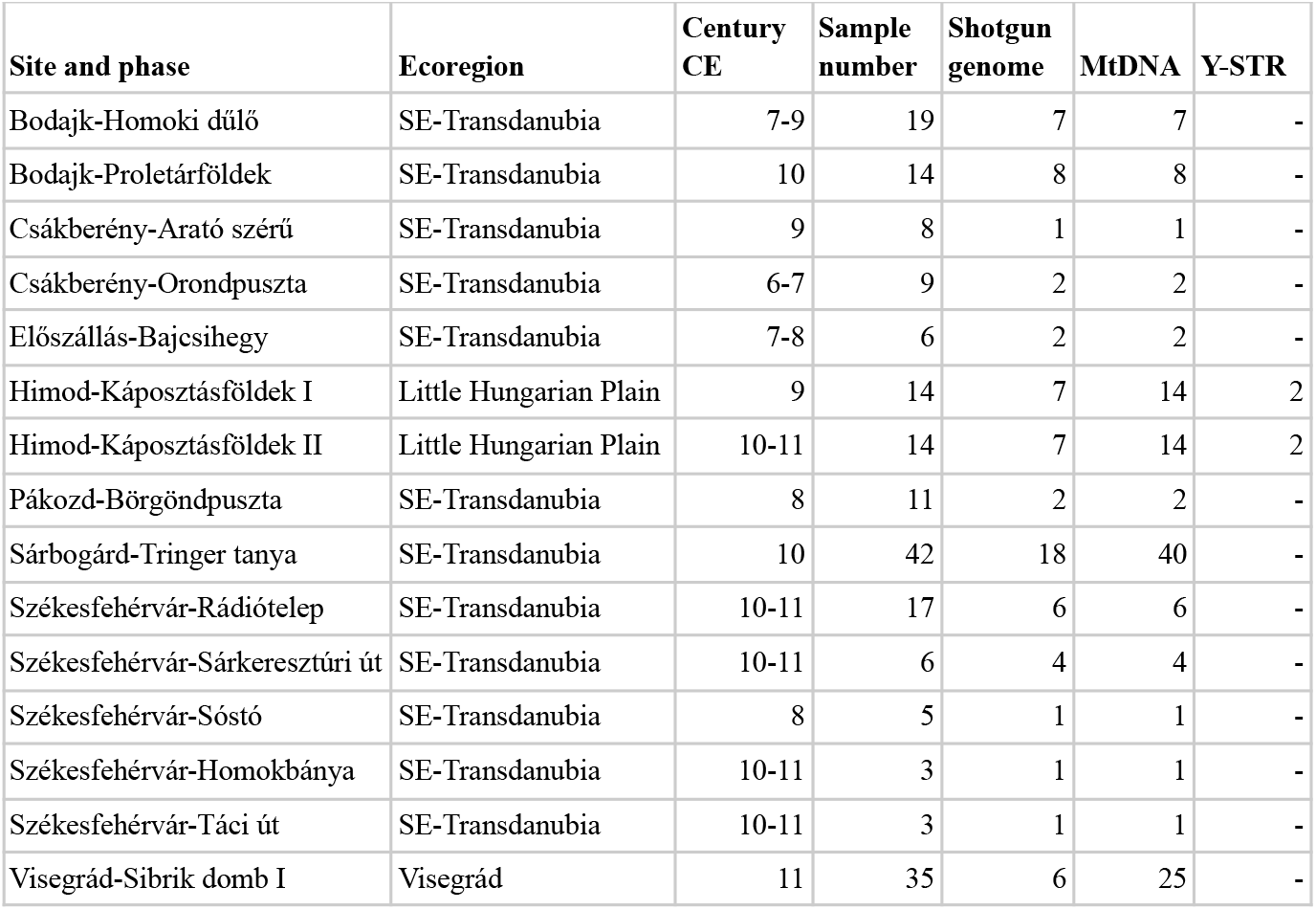
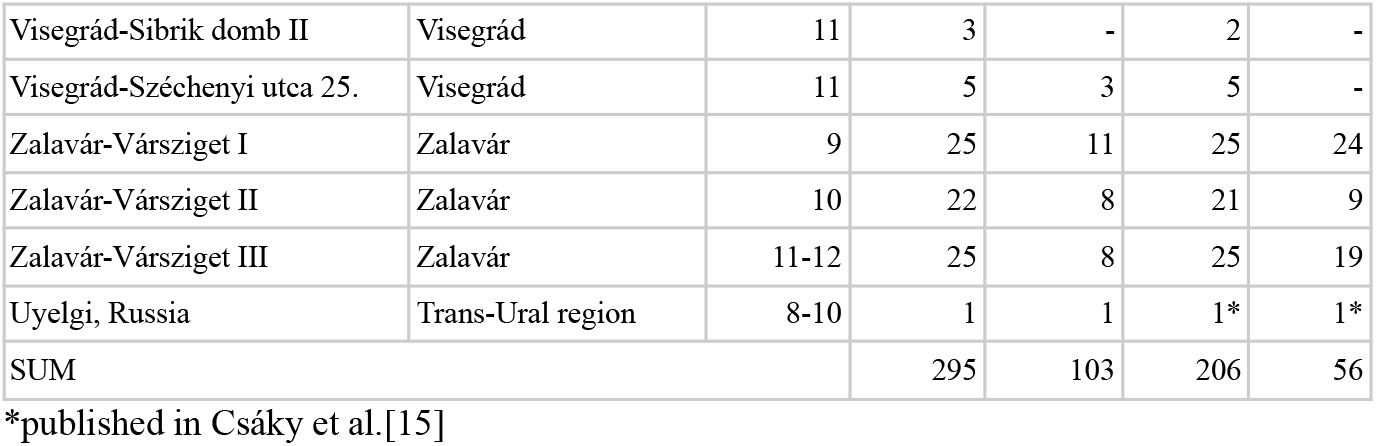
Summary of the sampling and analyses carried out in this study.

**Figure 1.**
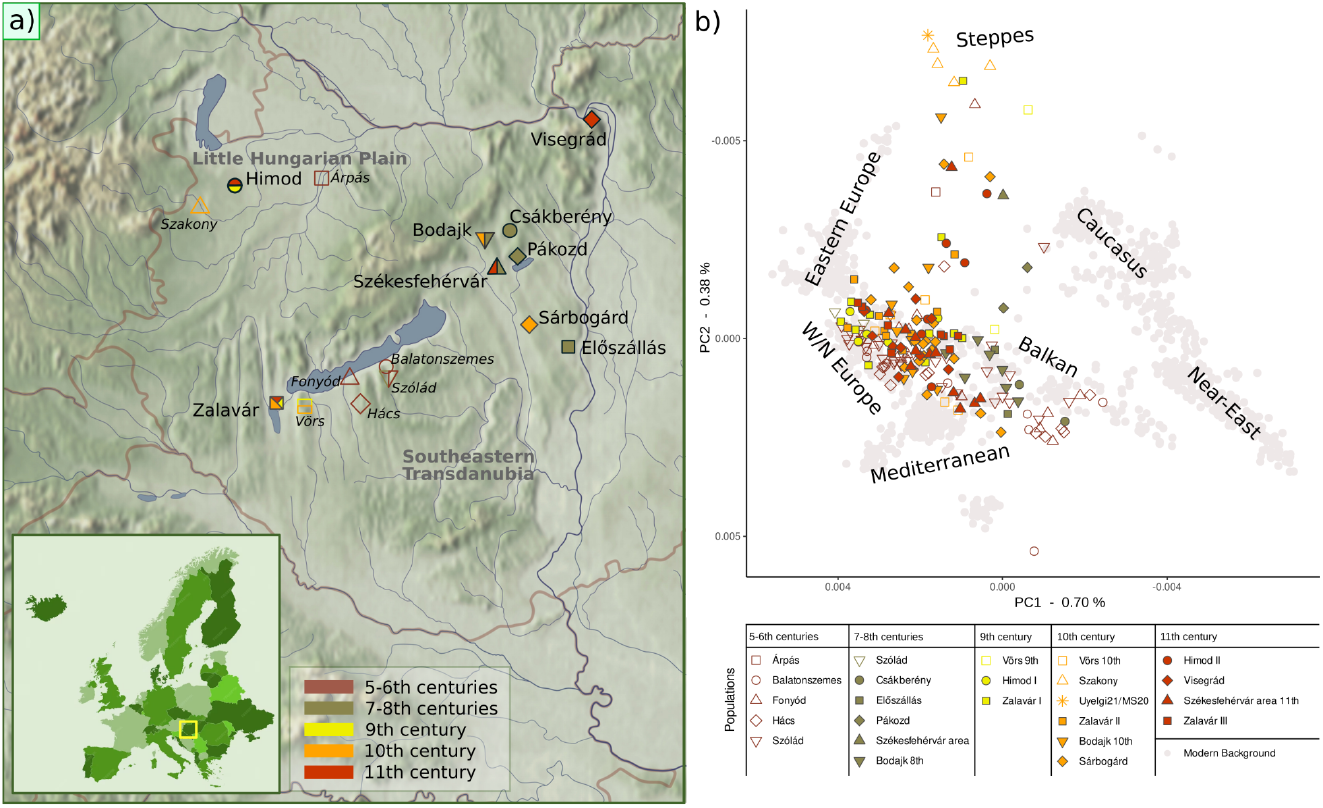
Sites and general genomic composition of the samples originating from Transdanubia. a) The discussed sites in Transdanubia, where italic labels and empty symbols indicate previously published datasets[8], [10], [11]. b) Principal component analysis (PCA) presenting the Transdanubian genomes from the 5th to 11th centuries CE. Ancient published (empty symbols) and newly sequenced (filled symbols) genomes are projected onto modern West Eurasian genomes (grey dots).

## Results

Out of the initially collected 295 human DNA samples, 103 were eligible for deeper shotgun sequencing, resulting in genomes with ∼0.019-2.14× average genomic coverage (mean ∼0.86×). The number of detected SNPs from the 1240k panel[14] ranges from 18,875 to 1,008,685 (mean ∼563,152), enabling a deep analysis of the dataset. In addition, 184 mitochondrial genomes via mitogenome capture and 56 Y-chromosomal STR profiles were created for samples from the Zalavár, Himod, Visegrád and Sárbogárd sites (Supplementary Information section 2, Table S1). Sample Uyelgi21, under the ID MS20 from the 8th-10th century CE Trans-Uralic Uyelgi site[15], was resequenced for its importance in Hungarian prehistory to 1.89× average genomic coverage (952,072 genomic SNP on 1240k, Table S1), and used as a proxy to distinguish Northeast-Eurasian and Western-Siberian genetic ancestries and connection systems in our analyses. For further sample information, see Supplementary section S1. In the following sections, we co-analyzed our dataset with aDNA samples from the region of present-day Hungary, dated between the 5th and 11th centuries CE[8], [9], [10], [11]. For *f*-statistics, we also included samples from Antonio et al.[16]. Accordingly, the complete dataset that was also eligible for IBD (identity-by-descent) analysis consisted of a final set of 416 genomes (Table S5).

### Genomic composition in the 5th-11th centuries CE Carpathian Basin

Both the principal component analysis (PCA) plot, projected onto a modern West-Eurasian background (Figures 1.b and S6-S10), and the K=6 unsupervised *ADMIXTURE* analysis (Fig. S11, Table S2) show large diversity among the studied samples from Transdanubia (5th-12th centuries CE). This diversity is also reflected in the uniparental makeup that represents a wide range of Eurasian macrohaplogroups at both the mitochondrial and Y-chromosomal levels (Supplementary Information section 2).

To describe the variability of the investigated dataset, we first separated individuals bearing only European ancestry through several *f*_4_ analyses[17] and tested these for population structure, further described in Supplementary Information section 3.2. Our analyses revealed three major genetic groups (1-3) and two subgroups of Group 3 (3.1 and 3.2) among individuals with only European ancestry. This is also reflected by PCA positions (Fig. 2a) and *ADMIXTURE* proportions (Fig. S13). After performing further *f*_4_ tests (Supplementary Information section 3.2), we found that these groups form a genetic cline in the region, distributed roughly in a North-South axis (Fig. 2a, Table S5). Groups 1 and 2 represent the extremes, while Group 3 is generally a mixture or cline between them. Therefore, we refer to it as European Cline 3 in the following analyses. This description is a refined version of the Carpathian Basin European diversity characterized by Maróti et al.[10]. We also discovered that Near-Eastern ancestry is strongly correlated with Group 1 and Cline 3.1 (Supplementary Information section 3.2). The genomic makeup of Group 1 and Cline 3.1, along with their PCA positions and uniparental markers, indicates a strong Late Antique and/or Balkan influence in the studied region, as also observed by Olalde et al.[18]. This suggests similar impacts in the territory of present-day Hungary compared to the Balkans during the first half of the first millennium CE. Similarly, Group 2 can be paralleled with one or more northern European impacts, as partially described by Vyas et al.[11]. Our *f*_4_ grouping results were also confirmed by subsequent IBD analyses (Supplementary Information section 3.2, Fig. S21). We refer to individuals with exclusively European, Near-Eastern, and Caucasian ancestry as CB-EUR hereafter, as the latter two often co-occur and do not show specific ingroup separation in regional IBD analyses (see below). Based on Olalde et al.[18], these ancestries appeared in the Balkans by the end of the Roman period and are considered to be part of the autochthonous population makeup of the studied era.

**Figure 2.**
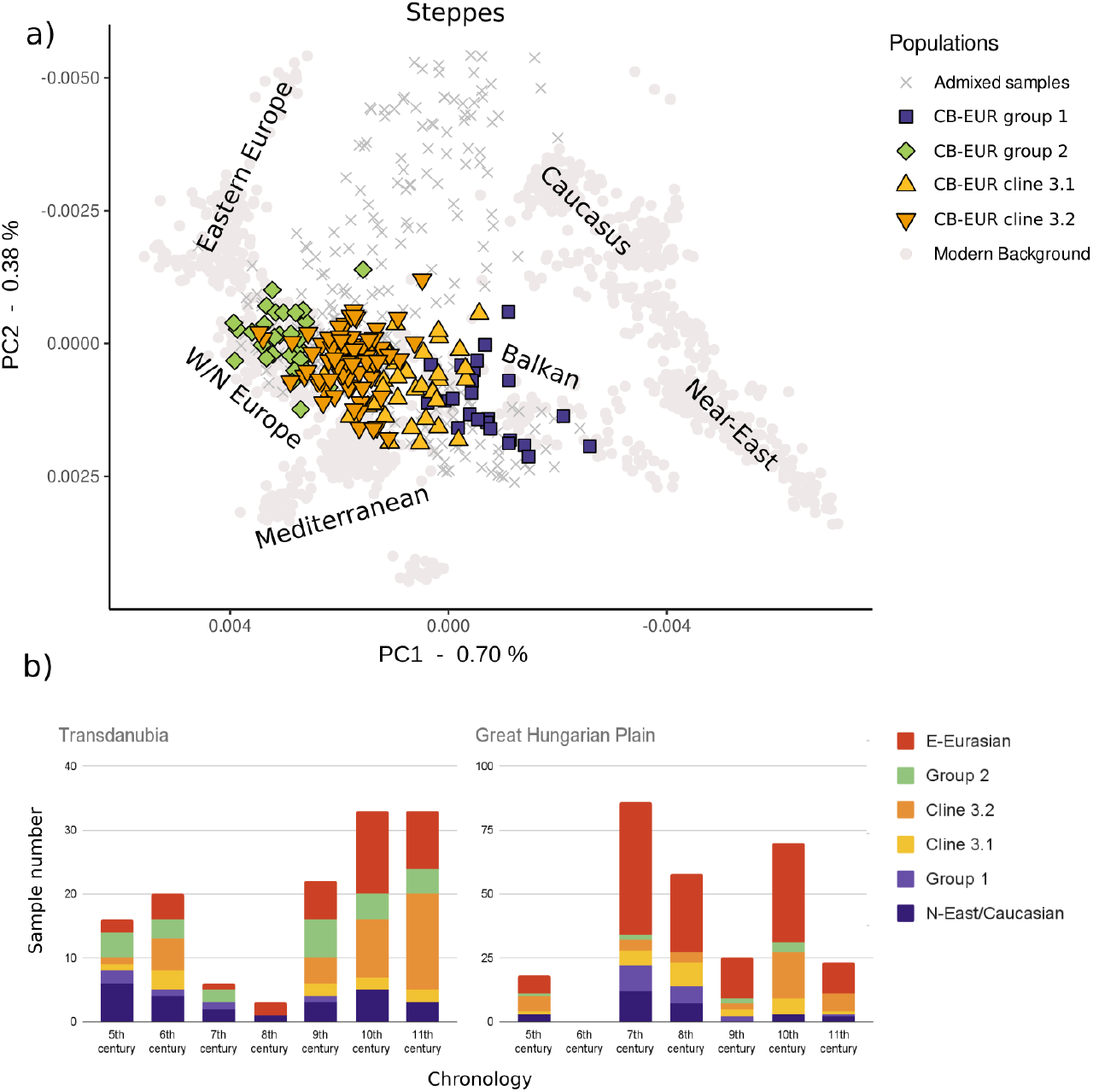
Distribution of European genetic groups. a) Principal component analysis (PCA) representation of f_4_-cluster based classification of individuals bearing only European ancestry. Further tests (Supplementary Information section 3.2) revealed that these groups represent sections of a regional genetic cline, where Groups 1 and 2 represent the two extremes, and Cline 3 the genetic cline between these. b) Chronological and geographical distribution of the genetic groups/clines. Until a homogenisation process in the 10th century CE, clear differences between the two macroregions can be observed.

The chronological and geographical distribution (Fig. 2b) of these genetic groups/clines and individuals with non-European ancestry varies over time and between Transdanubia and the GHP until the 10th century CE (Supplementary Information section 3.2, Fig. S16). Roughly half of all investigated samples possessed some non-European ancestry (tested by *f*_4_-statistics), with the prevalence of East-Eurasian ancestry being twice as high in the GHP compared to Transdanubia (Fig. 2b, Table S4). It is important to note that *f*_4_-statistics may not detect trace amounts of ancestries. Therefore, minor East-Eurasian ancestry may be present in the CB-EUR individuals, but this did not distort further analyses.

### Genomic haplotype segment sharing within the Carpathian Basin

Due to the methodological limitation of the allele frequency based methods[19], we also applied IBD analysis using *ancIBD*[20] on high-quality samples after *GLIMPSE* phasing and imputation[21]. For details, see Materials and Methods section, and for the results, see Supplementary Information section 4 and Table S6.

Text box 1: Benefits and drawbacks of IBD-based analyses.

1) The accuracy of phasing and imputation, which highly depends on the background database, genome coverage, and data type (capture or shotgun), significantly affects the correct recovery of shared IBD segments according to estimates by Ringbauer et al.[20]. Although ancIBD robustly detects IBD longer than 8 cM[20], we restricted analyses to >10 cM segments in order to minimize false signals.
2) Unlike allele-frequency-based population genetic analyses, the IBD-based analyses do not inherently require the exclusion of close relatives. However, for certain IBD-based analyses (as described in the Methods section), we opted to exclude relatives to avoid potential distortions in the results.
3) Shared IBD segments tend to shorten, and 10 cM segments disappear over a few hundred years, which is a phenomenon that must be considered when interpreting the data. Therefore, the lack of shared IBD segments between two individuals or clusters/groups, especially with a wider chronological gap, does not necessarily indicate a lack of genetic connections, but could still reflect no or limited shared genetic ancestry. Additionally, effective population size, community organization and sample number also influence the pattern of observed IBD sharing.
4) Recovering shared IBD segments between individuals and clusters/groups does not directly reflect the ancestry composition of the studied populations. Therefore this evidence needs to be discussed separately. Shared IBD segments between two individuals could either reflect a direct relationship between them or heritage from their common ancestry.

We defined degree centralities (corresponding to the number of IBD connections with sum IBD>10 cM, both individually and group-based), which vary across centuries and regions. Applying an edge-weighted spring embedded layout on the undirected graph, two major clusters are formed (Fig. S19) by individuals with predominant East-Eurasian ancestry. This largely overlaps with chronology – with the clusters separated by the event of the Hungarian Conquest approximately during the 890’s CE. The individual Uyelgi21, associated with the late Kushnarenkovo culture in the Southern Urals (8th-10th centuries CE), shows extensive connections to post-Conquest individuals with East-Eurasian ancestry (post-CEE) and only one 10 cM connection to a pre-Conquest cluster of individuals with East-Eurasian ancestry (pre-CEE), reinforcing the separation of the two clusters (Fig. 3). Individuals of CB-EUR exhibit a rather homogeneously scattered network, likely due to a larger population size (see below). Next, we redefined these East-Eurasian ancestry clusters based on their chronological distribution (pre-CEE: 7th-9th centuries CE, post-CEE: 10th-11th centuries CE). While the chronology and IBD cluster classification of East-Eurasian ancestry individuals generally align, we identified some exceptions (Table S5, detailed in the Discussion).

**Figure 3.**
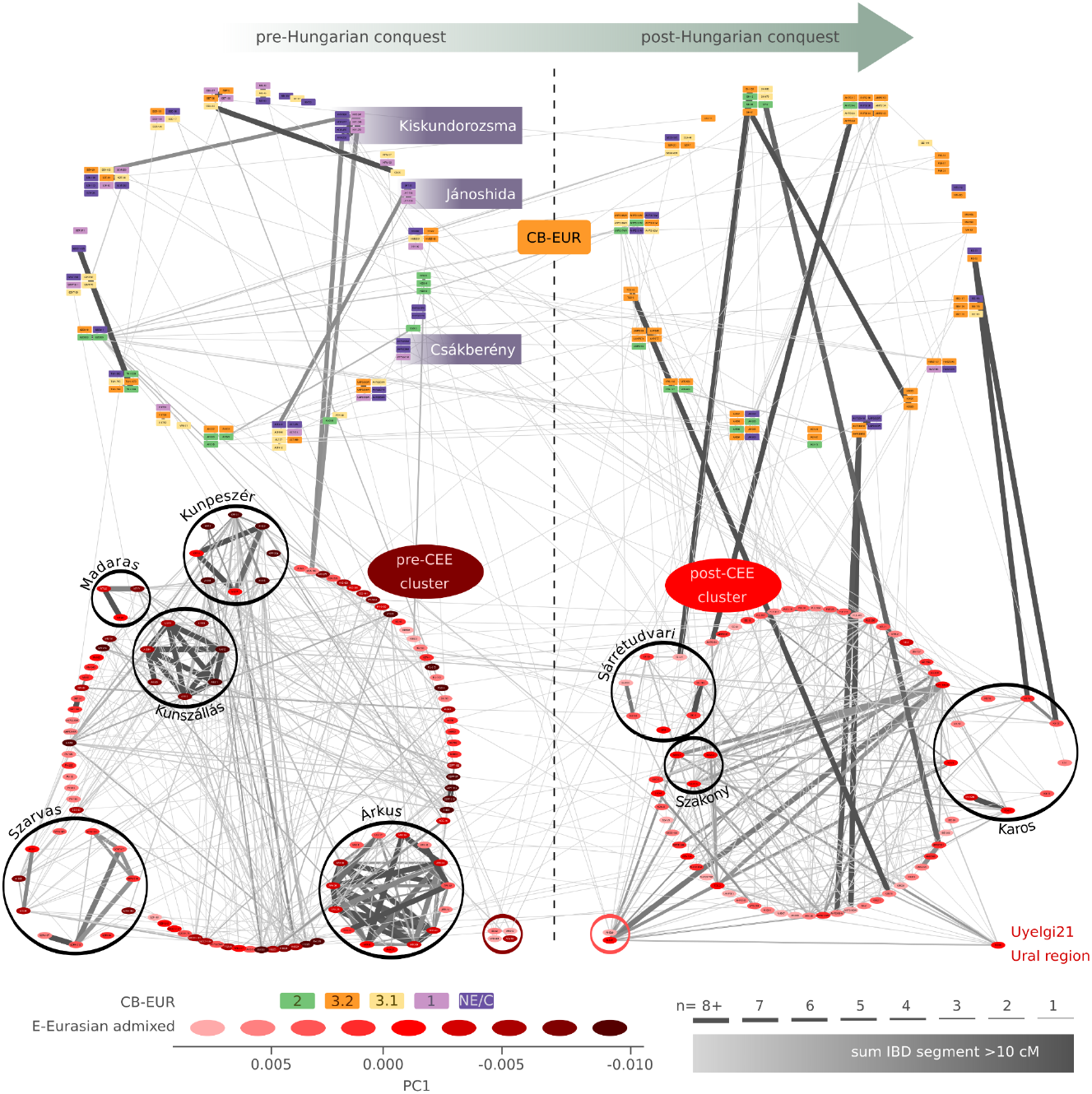
IBD (identity-by-descent) segment sharing networks of the Carpathian Basin populations in the 7th-11th centuries CE. The colouration of the individuals in CB-EUR corresponds to their European genetic groups/clines. The relative positions of the first principal component (PC1) are illustrated at the bottom. The samples are grouped by sites (Table S5). Ovals in the lower clusters represent individuals with significant East-Eurasian ancestry and color depth reflects their PC1 positions. Samples are sorted by sites, with those mentioned in the text circled in black. The width of the edges corresponds to the number of >10 cM IBD segments and the color depth signals the total shared length in cM (centimorgan). Three sites are highlighted for their homogeneous CB-EUR grouping. Individuals of different chronological and East-Eurasian cluster classification are separated and circled in red at the bottom of the graph. For individual classifications, see Table S5, for IBD connections, refer to Table S6.

Most of the post-CEE cluster individuals are situated on a different cline on the Eurasian PCA compared to the pre-CEE cluster individuals. Many are positioned towards and almost overlapping with Uyelgi21 (Fig. S12), consistent with its extensive IBD connections to them. Despite several 10-20 cM connections (Table S6.1), we find predominant discontinuity between pre- and post-CEE clusters as well (Fig. 3). Since the majority of individuals in the pre- and post-CEE clusters also possess West-Eurasian ancestry, we identified those completely lacking this component in Supplementary Information Section 3.2 (Table S4.3). This analysis aimed to determine whether the sporadic connections between pre- and post-CEE clusters are through CB-EUR individuals, local East-Eurasian admixture, or common East-Eurasian ancestry. Accordingly, we observed the latter only through a few connections (Table S6.1). These connections accumulate among post-CEE individuals (with one exception, see Discussion) from the second half of the 10th century CE in some scattered areas, indicating sporadic and regionally varied survival of these lineages into the post-CEE cluster, suggesting local admixture rather than common eastern ancestry.

The IBD connections of the pre-CEE cluster to CB-EUR are much more sparse compared to those of the post-CEE cluster (the average individual connection to CB-EUR for pre-CEE is 0.54 and for post-CEE is 1.39, see Fig. 3). Conversely, the average within-cluster connections for pre-CEE (∼3.16) and post-CEE (∼2.55) are rather similar, and are much higher, compared to the pre-CB-EUR (contemporaneous CB-EUR individuals to pre-CEE, ∼0.38) and post-CB-EUR (∼0.30) (for individual classification, see Table S5). The kinship level (1st-5th degree) connection network is remarkably different between the Avar and Conquest periods. Such connections of the pre-CEE cluster remain within-cluster, except for one sample (KDA-188, Kiskundorozsma site), and are all restricted to the same site. Conversely, while the post-CEE cluster connections are also mostly restricted to the same site, they primarily occur between CB-EUR and post-CEE individuals. Interestingly, both post-CEE males and females were involved in this admixture with CB-EUR individuals, likely in equal proportion. This conclusion is based on the distribution of presumably Eastern Y chromosomal (C2, Q1, N1a, R1a-Z93) and mtDNA haplogroups (A, B, C, D, G, M, N1a1a1a1a, N9, R, U4d2, X2f) found in the post-CEE individuals, alongside West-Eurasian uniparental markers (Table S10)[10], [15], [22]. Another striking feature is the relative isolation of the pre-CEE cluster: despite extensive admixture with West-Eurasians (based on PC1 positions), the paucity of CB-EUR connections suggests a non-local source for this component, consistent with the observations of Gnecchi-Ruscone et al.[23] and our *f*_4_-results (Supplementary Information section 3.2 and Fig. S16).

The IBD results, particularly the above outlined network density differences between clusters, become more conspicuous when considering the effective population size (*N*_*e*_) of the pre-CEE cluster compared to the post-CEE cluster, calculated with *hapROH* (see Methods). We calculated and compared the *N*_*e*_ of the pre- and post-CEE clusters to their contemporaneous subset of CB-EUR individuals (Table S8). Estimates on actual population size are problematic due to the lack of solid historical or anthropological demographic data for the period, but approximations were made nonetheless[24]. Our analysis suggests that around 20.82% of the population of the Avar period belonged to the pre-CEE cluster, whereas only approximately 13.01% of the population from the 10th-11th centuries CE belonged to the post-CEE cluster. This explains the IBD network density differences between the clusters (*i*.*e*. a smaller population has more internal IBD sharing), and aligns with historical estimates of 500k to 2 million individuals for the Carpathian Basin and 50k to 150k individuals for Hungarian conquerors (which may overlap with post-CEE, see Discussion)[24]. Additionally, the *N*_*e*_ estimates for CB-EUR indicate a small but definite increase towards the end of the millennium (Table S8). We note that *N*_*e*_ might be heavily influenced by societal customs and marriage patterns, therefore it is not necessarily indicative for census population sizes.

The IBD connections of CB-EUR (Fig. S21) reflect the *f*_4_-based classification and also suggest that different sections of the CB-EUR cline likely consist of distinct communities and groups. For example, those belonging to European Group 2 show discontinuity before and after the 7th century CE (Fig. S22), indicating the presence of different groups that share a similar regional genetic makeup.

### Population genetic events in the 5th-11th centuries CE

To further understand the social and regional dynamics of communities between the 5th and 11th centuries CE, we analyzed trends in community networks based on IBD data. We explored weighted degree centrality (i.e., the number of connections, denoted as k, calculated separately for between-site (k_B_) and within-site (k_W_) interactions) using different weighting methods (see Methods for details and Table S6.2 and Fig. 4 for results). We calculated *k*_*B*_ and *k*_*W*_ values separately for Transdanubia and the GHP, as well as for the combination of these two macroregions, denoted as ‘Merged’ in Fig. 4. The *k* values were also computed for each time interval (*i*.*e*. centuries), as shown in Figures 4c and 4d. This approach allows us to capture both spatial and temporal differences in the network structure. Furthermore, we were particularly interested in *k* values of the CB-EUR cluster alone, as it might behave differently compared to the broader dataset where non-local elements are overrepresented. Due to the low number of CB-EUR connections, we developed a cumulative approach for *k*_*B*_. Although we used a different formula with two separate weighting methods (see Methods), the theory is similar to the conventional approach, as it involves using a weighted number of connections and adding samples from previous batches to each subsequent batch. The rationale behind this approach is to detect connections that might otherwise be masked by the narrow time window. Using our formula, this method highlights local trends that would otherwise remain hidden (Figures 4a and b).

**Figure 4.**
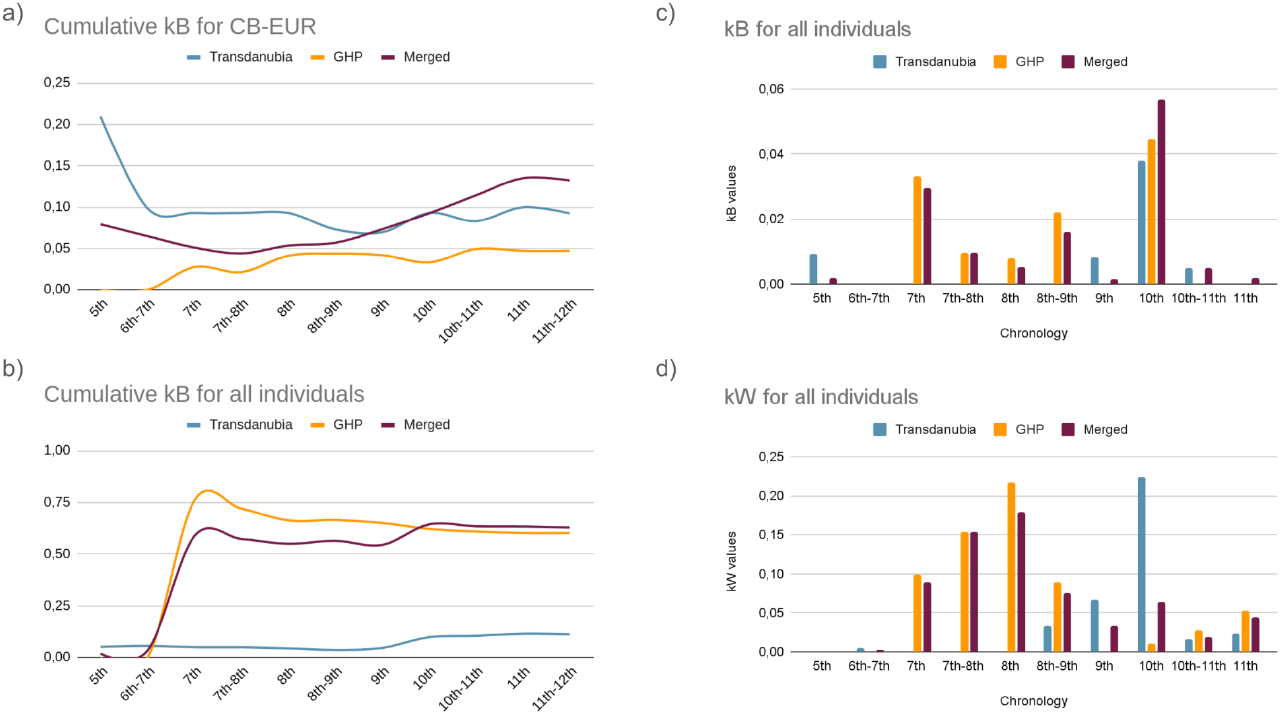
Intra- and intergroup tendencies in IBD (identity-by-descent) segment sharing patterns. a) Cumulative connections between sites (k_B_) regarding the CB-EUR cluster in a chronological order, indicate movement into and within Transdanubia during the 10th century CE but not within the GHP (Great Hungarian Plain). b) The same analysis, when including all individuals (also CEE), signals contact between sites in the GHP during the 7th century CE, which likely can be connected to the settlement and exogamous social structure of the Avars. c) The non-cumulative approach of connections between sites signals the previously observed two waves of movements. d) Within site connections (k_W_) reveal moderate biological networks across sites (Table S6.2).

Cumulative *k*_*B*_ trends differ remarkably when calculating these for only CB-EUR versus all individuals, including those with East-Eurasian ancestry (Figures 4a and 4b), which points to vastly different social organizations of local (CB-EUR) communities compared to those who possess East-Eurasian ancestry. However, certain commonalities also emerge: 1) The ‘Merged’ values, which signify connection between the Transdanubian and the GHP communities, shows the same relative trend in both cases. Accordingly, Transdanubia and the GHP remained relatively separated until the turn of the 9th-10th century CE, when, likely fueled by the Hungarian conquest, contacts between these regions increased. 2) The GHP communities, contrary to the Transdanubian communities, show no change in *k*_*B*_ values during this turning point, pointing to movements from the GHP to Transdanubia. This trend is also much more pronounced for the Transdanubian CB-EUR group when using one of our alternate weighting methods (Methods, Fig. S23). The sudden increase of intersite connections in the 7th century CE onwards in the GHP communities are in line with the observations of Gnecchi-Ruscone et al.[23], who suggested sedentary lifestyle along with female exogamy within the eastern-origin strata of the Avar period.

The conventionally calculated *k*_*B*_ (Fig. 4c) revealed two major peaks in the 7th and 10th centuries, indicating two waves of intensive biological contact between sites, and stability in between. The first wave correlates with the changing Avar society in the 7th century CE, while the second wave aligns with the reorganizations fueled by the Hungarian conquest at the turn of the 9th-10th centuries CE. The stable lifestyle in rather smaller communities is reflected by some 7th century CE sites, such as Kunpeszér and Kunszállás from Maróti et al.[10] and Gnecchi-Ruscone et al.[9], and even by the few samples from Madaras (Fig. 3), a large burial site of the 7th century CE, which show high degree of within-site connections, regardless of biological sex. This observation further diversifies previous findings about female exogamy during this period[23]. In addition to that, the data also show a notable rise of between-site connections at the turn of the 8th-9th century CE in the GHP, indicating some degree of population movement or change during that time. In Fig. 4d, the *k*_*W*_ values complement the *k*_*B*_ in Fig. 4c. There are several notable features in the joint interpretation of Figures 4c and 4d. The first is the increased contacts around the turn of the 8th-9th centuries CE in the GHP compared to the previous century, which may reflect events related to the changing political environment described in the introduction, although other scenarios are also possible. Another notable feature is the high *k*_*W*_ value for 10th century CE Transdanubia, especially in comparison to that of the GHP. This phenomenon could result from chronological assignment bias, although the sample number is the highest in this 10th century CE batch (103 overall, 33 for Transdanubia), and the *k*_*B*_ values of the same batch are synchronous with those of the GHP, making this scenario unlikely. If the observed *k*_*W*_ is a true feature of the dataset, it may signal a few generations’ delay of movements in Transdanubia compared to the GHP. This scenario correlates with archaeological-historical data that the Hungarian conquest was a sequence of events in the Carpathian Basin from the eastern to western territories[25]. On the other hand, this phenomenon could also be the result of different sampling strategies between the Transdanubian and GHP sites, as in the study by Maróti et al.[10] many characteristically military-related cemeteries were sampled, which have a different organization and IBD pattern compared to biological community-based community burial sites. After the Hungarian conquest, *k*_*W*_ values tend to slowly increase in the following centuries in both regions, pointing to a more stable and established community system.

## Discussion

### Fluctuations within the CB-EUR cluster throughout the 5th-10th centuries CE

The chronologically changing makeup of distant segments of the CB-EUR genetic cline defined by our analyses reveals complex population processes in the Carpathian Basin during the second half of the first millennium CE. European Group 2 individuals are rather abundant in Transdanubia, along with related Cline 3.2. A common element between regions is the proportional decline of European genetic Group 1 and Near-Eastern ancestry, as well as the proportional enrichment of Cline 3 (*i*.*e*. the integration of Groups 1 and 2 into Cline 3) towards the 11th century CE. The proportional similarities of the two regions in the 10th century CE suggests an amalgamation process of different groups across the studied area, which we confirmed by IBD segment connection dynamics as well.

We highlight that the presence of clinal similarity between different regions or horizons does not necessarily indicate group identity, even at a biological level, as European Group 2 shows discontinuity pre- and post-7th century CE based on IBD results (Fig. S22). This observation underscores that PCA and evolutionary-oriented techniques, such as *ADMIXTURE* and *f*-statistics based methods, are limited in their ability to correctly characterize populations in a frequently intermixing milieu. Therefore, conclusions drawn from such results must be approached with great caution. However, certain trends should not be overlooked. For example, in the pre-Conquest period, the presence of a remnant Romanised population around Csákberény, based on archaeological findings (Supplementary Information section 1.8), is paralleled by the Group 1/Near Eastern genetic makeup of the individuals from this site (Fig. 3, Supplementary Information section 3, Results). The accumulation of certain elements further suggests some local isolation in Himod I with Group 2 individuals, and the GHP sites Kiskundorozsma and Jánoshida[10] with Group 1 individuals, although these are less corroborated by archaeological background (Fig. 3). This genetic isolation is, however, mitigated by connections to pre-CEE individuals, such as one in Kiskundorozsma.

### Dynamics of the communities with East-Eurasian ancestry before and after the Hungarian Conquest period

Recent researches have highlighted the predominance of East-Eurasian ancestry among the early Avar Khaganate’s elite, indicating these groups as a significant genetic source of immigrants (Fig. S12)[9], [10], [23]. Therefore, we conclude that the pre-CEE cluster defined in this study mostly represent the East-Eurasian core and genetically related individuals of the Avars, although we acknowledge that the ethnic identities and genetic categories used do not correspond directly. Similarly, the post-CEE cluster can be strongly associated with the Hungarian conquerors and their descendants. The remarkable discontinuity between pre- and post-CEE clusters is nuanced by the sporadic IBD connections, which almost exclusively accumulate in cemeteries from the second half of the 10th century CE (Table S6.1), indicating limited local survival of pre-CEE cluster lineages. The only systematic exception is Karos cemetery, which represents one of the earliest Hungarian conqueror cemeteries in the Carpathian Basin but also shows the strongest IBD connections to the pre-CEE cluster. On the other hand, the diverse genetic backgrounds observable here indicate the varied origins of this community’s members, framed within a military unit, as evidenced by the ratio of armed male burials and the diverse cultural elements found in the cemetery material[26]. Additionally, contrary to the suggested continuity of 5th century CE Hun period East-Eurasian genetic ancestry into the Conqueror cluster by Maróti et al.[10], our data indicate an absence or loss of this ancestry among post-CEE individuals in the studied region (Fig. S24). However, the greater time gap and the limited number of Hun period samples in the Carpathian Basin may obscure distant and sporadic genetic connections.

The IBD connections of the Hungarian conquerors to CB-EUR demonstrate an immediate mingling with the locals, while the rarity of such connections between the eastern Avars and the parallel CB-EUR (Fig. 3) is striking, especially considering these groups relative population size. This difference is also highlighted by the results of *f*_4_-statistics, which suggest different West-Eurasian ancestry for pre- and post-CEE clusters (Supplementary Information section 3.2, Fig. S16). These results, however, can be interpreted in various ways. The genetic isolation of the Avar elite previously demonstrated by studies[9], [12], [23] is now corroborated by our IBD analyses. This highlights the distinct social organizations of the communities belonging to the pre- and post-CEE clusters. Our findings provide a nuanced perspective on the observations of Gnecchi-Ruscone et al.[23], regarding extended exogamy as a general residence pattern of the Avar society. Based on the results summarized in Fig. 4, Avar communities show two waves of between-site connections (likely driven by female exogamy) first in the 7th century CE and again at the turn of the 8th-9th century CE. This observation is further supported by the high within-site kinship-level connections of late Avar period sites, such as Árkus and Szarvas (Fig. 3), irrespective of biological sex. Notably, at the sites investigated by Gnecchi-Ruscone et al.[23], many females did not connect to the studied pedigrees, and many pedigree-forming females are missing from the family trees. In a broader multi-site (macroregional) model, our findings depict a different social structure. However, we cannot rule out exogamy from closely neighboring villages during periods where we observe a drop in *k*_*B*_ values. This idea is also supported by the strontium isotope results from Gnecchi-Ruscone et al.[23], which did not indicate large-scale movement between distant villages. The relative isolation found among the Avars (Fig. 3) may explain their eventual disappearance and the subsequent genetic and cultural succession by the Hungarians. Furthermore, these results suggest that the West-Eurasian (likely Eastern-European, Steppe and/or Caucasian) admixture in the pre-CEE cluster primarily originated outside the Carpathian Basin, consistent with previous observations[23]. This indicates a prevailing West-Eurasian genetic substrate during the Avar period, which is currently traceable only through its admixture into the elite East-Eurasian families.

### Microregional genetic trends in Transdanubia

Focusing on the IBD network of Transdanubia, we grouped sites according to their ecological and chronological distribution (Fig. 5, Supplementary Information section 4). Two individuals from the early 7th century occupation of Szólád (AV1 and AV2, first-degree relatives) display extensive connections to individuals from most of the studied sites, with the most notable connections being to Zalavár (*Mosaburg*). Recovered from deposit fillings of a Langobard period grave without any artifacts[8], these individuals nonetheless indicate the presence of a genetically successful (and likely large) substrate in the region, probably arriving around the 7th century CE. Given this genetic substrate’s presence centuries before *Mosaburg* was founded, we cannot confirm the written sources that claim the entire settlement’s population originated outside the Carpathian Basin, suggesting possible exaggerations in historical records. Nevertheless, the presence of mtDNA haplogroup C1e (AHP20), previously only found in Icelandic populations[27], and the Y-chromosomal haplogroups I1-Y6357 (AHP16) and R1a-FGC11904 (AHP18), primarily of Northwest-European origin (Yfull.org) suggest some foreign connections in the Carolingian horizon of Zalavár. Although the sample size is small relative to the presumed large populations of the past, the Zalavár sample set demonstrates some continuity from the 9th to the 11th centuries CE.

**Figure 5.**
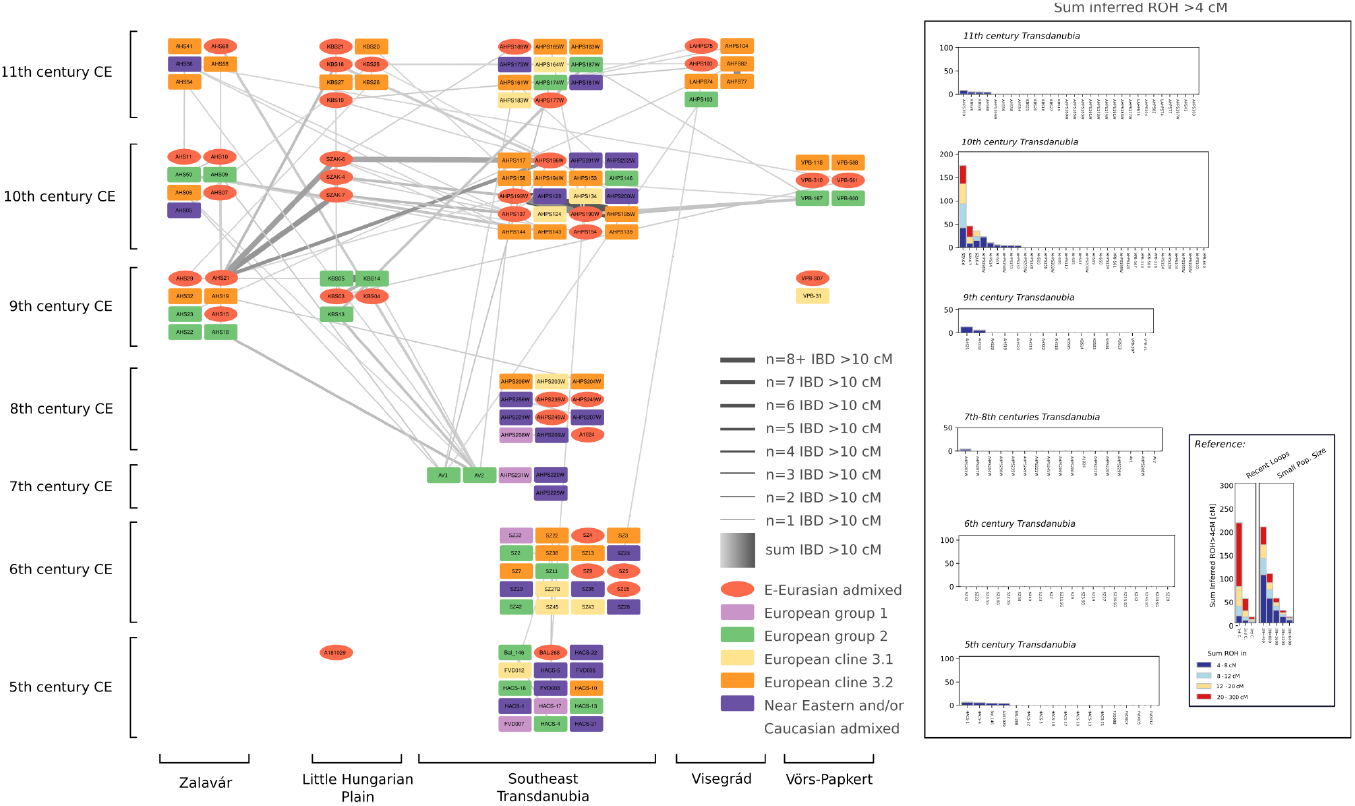
IBD (identity-by-descent) patterns across Transdanubian sites in the 5th-11th centuries and their corresponding ROH patterns. Only connections between 10 and 1400 cM (centimorgan) are visualized, in order to emphasize population level connections instead of 1-2nd degree connections.

The 9th century CE phase of the northwestern Transdanubian Himod site also shows connections to AV1-2 (7th century CE, Szólád site, Transdanubia) and even the accumulation of CB-EUR Group 2 elements. Connections between the pre- and post-Conquest phases (I-II) of the Himod site are sporadic, and differences in admixture proportions (Fig. S11) and uniparental makeup suggest limited genetic continuity between the horizons. Instead, these results rather reflect new Hungarian occupation, with incoming new genetic components connected to the GHP and the post-CEE cluster, and a subsequent admixture process in Himod II. The northeastern Transdanubian Visegrád site shows more IBD connections to contemporary groups rather than preceding ones in other regions, corroborating written sources that describe it as a newly formed community[28]. However, future comparative IBD analyses with other European areas are required to fully understand the composition of this society. In the thoroughly analyzed Székesfehérvár area, we observe a different pattern compared to Zalavár. A Late Antique-type population persisted in this region, and could even be found around Sárbogárd in the 10th century CE. However, the appearance of Eastern elements after the Hungarian conquest is much more prominent in the area. Despite the larger number of samples from southeastern Transdanubia, signs of local succession are less pronounced in this area. This is consistent with expectations of community restructuring after the Hungarian conquest, as described by the IBD degree centrality changes throughout the centuries. Overall, the microregional evaluation reveals spatially different histories even between communities in close geographical proximity, highlighting the importance of prudently integrating archaeological and biological perspectives when interpreting such datasets.

### Changing communities in the late 9th - early 10th century Carpathian Basin

The observed changes in connections between and within the sites throughout the 5th-11th centuries reflect both Avar and Hungarian conqueror occupations and the amalgamation process of CB-EUR Groups 1 and 2 into Cline 3 towards the end of the millennium. Our analyses revealed that the GHP was genetically rather isolated from Transdanubia before the 10th century CE. However, the Hungarian conquest fueled intensive population movements and admixtures within the Carpathian Basin, consistent with historical records [5]. This process is further evidenced by the occurrence of relatives in geographically distant cemeteries after the Hungarian conquest. For instance, individuals KOZ018 (Himod, Transdanubia, 10th-11th century CE) and TCS-18 (Tiszanána, GHP, 10th century CE) are a father-son pair, while KOZ026 (Himod, Transdanubia, 10th-11th century CE) and SH-81 (Sárrétudvari, GHP, 10th century CE) are 3rd-degree relatives (Table S6). We observed a potential delay in this process in Transdanubia compared to the GHP, which underscores another difference between these two microregions, and highlights the importance of considering ecological barriers, such as the Danube River, in reconstructing past population events.

We also found evidence for the pre-conquest presence of individuals with Uralic ancestry. In phase ZVI (dated to 850-890), the adult male AHP21 shows strong connections to multiple individuals in the post-CEE cluster. This individual’s genomic composition is highly similar to Uyelgi21 with strong IBD connections (Table S6 and Fig. S12), and is a 5th-degree relative of adult male KeF1-10936 from Kenézlő-Fazekaszug (GHP), whose dating can only be approximated to 900-960 CE. Additionally, AHP29 from the Carolingian era (ZVI) also shares IBD segments with the Conqueror cluster, although its connections are much weaker compared to those of AHP21’s. Both burials firmly predate the Hungarian conquest based on archaeological stratigraphy (Supplementary Information section 1). These findings signal sporadic appearances of Hungarians in Transdanubia before 890 CE (new radiocarbon date of AHP21 is in Table S1, stratigraphic date is in Supplementary Information section 1). These individuals likely served as mercenaries at the court of Priwina, according to historical records (see Supplementary Information section 1.1).

### Limitations and final remarks of this study

Our results revealed complex population histories and both macro- and microregional events in the Carpathian Basin. We believe that the combination of the database data with ours, as well as the methods we used, provides enough statistical strength to the statements of this study. However, since we did not include comparative datasets from other countries—focusing instead on the local characterization of population events—we acknowledge that certain questions remain open. These include the ancestry shift within CB-EUR Group 2, the origins of the Late Antique-type population, and a more detailed characterization of the post-CEE cluster. These issues could be more fully addressed through exhaustive comparisons with data from other populations in future research. We believe that further sampling from East-Central Europe is required to reveal such intricate questions, and we aim to do so for subsequent studies.

## Materials and Methods

### Ancient DNA laboratory work

Samples were processed in a dedicated laboratory aDNA facility at the Institute of Archaeogenomics, Research Centre for the Humanities, HUN-REN. Sample preparation has taken place according to Szeifert et al.[22]. Bone powders were produced based on Lipson et al.[29] and Sirak et al.[30], DNA extractions were performed following the methods described in Dabney et al.[31] and Rohland et al.[32], and the double stranded DNA libraries were created using half-UDG treatment[33]. DNA extraction and library preparation were performed either manually or automatically with Biomek i5 Liquid Handling Workstations. Initial screenings were performed on an inhouse Illumina MiSeq platform, and eligible samples were then deeper sequenced on high throughput Illumina platforms (Applied Biotechnologies and Novogene), for all results, see Table S1.

We performed mtDNA hybridisation capture[22], [34] and Y chromosome STR analyses using AmpFlSTR® Yfiler® PCR Amplification Kit (Applied Biosystems). Y chromosome haplotype analysis was carried out in GeneMapper® ID Software v3.2.1 (Applied Biosystems).

### Bioinformatic analyses

The *PAPline* package with default settings[35] were used for Illumina paired-end sequencing reads mapping to *Homo sapiens* reference genome GRCH37.p13, pre- and post-filtering steps and mitochondrial sequences calls, for results, see Table S1. Y chromosome haplogroups were classified using the *Yleaf* v2.2 software[36]. Mitochondrial contamination ratio was estimated with *contamMix* (v1.0-11)[37].

The ratio of male X-chromosomal contamination was measured, with both *ANGSD* (version 0.939-10-g21ed01c, htslib 1.14-9-ge769401)[38] and *hapCon* (*hapROH* package version 0.60)[39]. The starting file for the latter method was a *samtools mpileup* (version 1.10, htslib 1.10.2-3ubuntu0.1) output with mapping quality 25 and base quality 30. All the parameters used were the default ones recommended in the official documentation. Similarly, the chrX_1240k.bed and chrX_1240k.hdf5 files used were supplied by the developers with the software package.

*ANGSD doCounts* was run on the X-chromosomal region from position 5500000 to 154000000, with mapping and base qualities 25 and 30 respectively. The resulting file was processed then with the toolkit’s contamination executable, with the following parameters: -b 5500000 -c 154000000 -d 3 -e 100 -p 1. The required HapMapChrX.gz file (-h parameter) was the one shipped with the software.

Kinship relations were estimated using *READ*[40] and *KIN*[41], and where available, supplemented by IBD connections (for results, see Table S2). For PCA, admixture analysis, and *f*-statistics, the dataset was merged with the AADR database v54[42]. PCA was calculated using the *smartpca* software[43] with the 590k SNP panel[42], for admixture analysis we used the *ADMIXTURE* software[44] with the 1240k SNP panel[42]. For *f*-statistics, we used the *Admixtools* software package[17], [45]. For the *ADMIXTURE* analysis, we used a dataset which contains genomic data of archaic age individuals from European, Near Eastern, Central Asian, and Mongolian populations. We filtered out the closely related individuals and those samples that were not treated with UDG.

We used *hapROH*[46] to detect runs of homozygosity (ROH) that indicate consanguinity of parents and small population sizes. The *Ne* vignette of *hapROH* was used to estimate effective population sizes (*N*_*e*_) with confidence intervals for *f*_4_ statistics-based genetic groups considering also chronological phases. 95% Confidence Intervals were fitted via both the likelihood profile (calculating the log likelihood for a large number of 2*N*_*e*_, and looking for the interval 1.92 log-likelihood units down from the Maximum Likelihood) and using the curvature of the likelihood (the so-called Fisher Information matrix). We also used different ROH segment ranges (4-20 cM, 4-8 cM, 8-12 cM, 12-20 cM) to monitor whether they show different temporal trends, as larger segments signal more recent inbreeding or population size reduction. The results are listed in Table S8.

Individuals with >250k SNPs on the 1240k panel were targeted for IBD analysis. Imputation was performed using *GLIMPSE* software[21], following the recommendations of the authors and using the v1.1.1 versions of the precompiled static binaries. The reference panel for the imputation (1000 Genomes Project (30X from NYGC) on hg19/GRcH37) was provided by Ali Akbari (Harvard Medical School). Genotype likelihoods of the samples were computed with a standard method involving bcftools *mpileup* and *bcftools call* v1.10.2[47]. Base- and mapping quality (flags *-Q* and *-q*) were both set to 20. As a difference from the official workflow, chromosomal level imputation regions were determined on a per sample basis, with the GLIMPSE_chunk_static executable, using default parameters (*--window-size 2000000 --buffer-size 200000*). The script used to phase and impute the samples is available on our github repository: https://github.com/ArchGenIn/AHP_Gerber2024/tree/main. To identify the IBD segments, we used *ancIBD* (version 0.5)[20] with the same process and parameters described in the software’s official documentation. The fraction of 1240k variants above 0.99 genotype probability on Chromosome 3 must have been at least 0.8 after imputation to include a sample for IBD calling. All IBD results were visualized using *Cytoscape*[48]. We restricted connections down to 1×10 cM, due to the high number of false-positive hits at shorter segments resulting from technical difficulties (*e*.*g*. phasing errors) and the sparse and asymmetric background database for imputation discussed further in the description of Fig. S25.

We also measured degree centrality (*k)*, using sum IBD connections between 10 and 1400 cM to exclude close relatives, as we were interested in the population-level processes. The conventional approach used for different time section batches (*i*.*e*. centuries) is the following for *k*_*W*_:

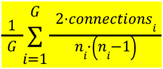

Where *G* is the number of groups within a batch, *connections*_*i*_ is the number of IBD connections within the i^th^ group (site) within the batch, and *n* is the number of individuals within the i^th^ group.

For *k*_*B*_ the following formula was used:

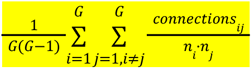

Where *connections*_*ij*_ is the number of IBD connections between the pair of the i^th^ and j^th^ sites. For the cumulative approach (applied for *k*_*B*_ only), we used:

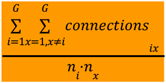

Where *connections*_*ix*_ denotes all IBD connections between the i^th^ and all other sites within the batch.

We also used an alternate weighting method to highlight global trends visualized in Fig. S23:

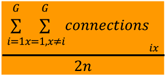

This latter normalization method balances sample size differences between sites and chronological batches, and thus highlights relative differences. The sample number distribution is summarized in Fig. S20. The script used for this calculation is also uploaded to our github repository.

## Abbreviations

CB-EUR: genetic cluster/cline in the Carpathian Basin of individuals with European, Caucasian and Near-Eastern ancestry;
GHP: Great Hungarian Plain;
IBD: Identity-by-descent;
*k*_*B*_: degree centrality of IBD connections between archaeological sites;
*k*_*W*_: degree centrality of IBD connections within archaeological sites;
post-CEE: post-Conquest East-Eurasian, genetic cluster of individuals with East-Eurasian ancestry chronologically following the Hungarian conquest;
pre-CEE: pre-Conquest East-Eurasian, genetic cluster of individuals with East-Eurasian ancestry chronologically preceding the Hungarian conquest;
SNP: Single nucleotide polymorphism;
STR: Short tandem repeat;
UDG: Uracil-dehydrogenase;
ZV: Zalavár-Vársziget site.

## Acknowledgement

Dr. Mihály Molnár (MTA ATOMKI HEKAL, AMS laboratory) is responsible for the radiocarbon analyses from Visegrád site.

On behalf of the ‘bioinfo-baden’ project we are grateful for the computational possibility to the HUN-REN Cloud[49] (https://science-cloud.hu/), which helped us achieve the results published in this paper.

We’d like to thank Ali Akbari for his help with the imputation of the whole genome data. We also thank Harald Ringbauer and Yilei Huang for their suggestions on running and optimizing ancIBD.

We are also grateful to the following archaeologists for their contribution to the project: Péter Tomka (for his advices about Himod-Káposztásföldek), Judit Kodolányi (Visegrád-Sibrik hill I), Gergely Buzás and Szabina Merva (Visegrád-Sibrik hill II), and István Kováts, Péter Gróf (Visegrád-Széchenyi street), Ágnes Ritoók (Zalavár-Vársziget), as well as anthropologist Tamás Szeniczey. We would like to thank for the invaluable comments on the paper to Péter Langó, Tivadar Vida, István Koncz, Balázs Gyuris and Levente Samu.

## Funding

The research presented in this article was funded within the framework of a subproject of the Árpád House Program, initiated by the Hungarian state (The Anthropological and Genetic Image of the Hungarians during the Árpád era, Árpád-house IV. framework program (1109/2018. (III.19.) Gov. decision).

Further funding of the research was provided by the Priority Research Theme proposal of the Eötvös Loránd Research Network (2019-2023 ELKH, 2023-HUN-REN), in the frame of the “Archaeogenomic research of the Etelköz region” project.

The radiocarbon investigations were carried out in the framework of the OTKA NK 104533 research project of Miklós Takács, which we are grateful for.

## Author contribution Ethics Statement

The authors declare that they had requested and received permission from the stakeholders, excavator, and processor anthropologists and archaeologists for the destructive ancient DNA analyses of the human remains presented in this study.

## Data availability

All data needed to evaluate the conclusions in the paper are present in the paper and/or the Supplementary Materials. The newly produced sequence data is deposited in the European Nucleotide Archive (ENA) with the following accession number: PRJEB75908. We also report on samples that have not yielded sufficient ancient DNA in Table S1.

