## Supplementary Information for "Ancient genomes reveal Avar-Hungarian transformations in the 9th-10th centuries CE Carpathian Basin"


**1 - Characteristics of the surveyed cemeteries and descriptions of the graves 30**
1.1 Zalavár-Vársziget 30
1.2 Himod-Káposztásföldek 38
1.3 Sárbogárd-Tringer tanya 40
1.4 Visegrád 44
1.4.1 Visegrád-Széchenyi street 25. 44
1.4.2 Visegrad-Sibrik Hill, site I 45
1.4.3 Visegrád-Sibrik Hill, site II 48
1.5 Sites of Székesfehérvár area 48
1.5.1 Székesfehérvár-Rádiótelep 48
1.5.2 Székesfehérvár-Sóstó, -Homokbánya, -Vízművek and -Sárkeresztúri út 49
1.5.3 Székesfehérvár-Táci út repülőtér (airport) 50
1.5.4 Székesfehérvár-Sóstó (Ikarus factory) 50
1.6 Bodajk 50
1.6.1 Bodajk-Proletárföldek 50
1.6.2 Bodajk-Homoki dűlő 51
1.7 Előszállás-Bajcsihegy (Mezőfalva-Vasútállomás) 53
1.8 Csákberény 53
1.8.1 Csákberény-Orondpuszta 53
1.8.2 Csákberény-Arató szérű 54
1.9 Pákozd-Börgöndpuszta (Székesfehérvár-Börgöndpuszta) 55
**2 - Uniparental analyses 56**
**3 - General genomic composition 57**
3.1 PCA and Admixture 57
3.2 f-statistics 57
**4 - IBD analyses 59**
**5 - Genetic relatedness analyses 60**
**Supplementary Figures 62**

### 1 - Characteristics of the surveyed cemeteries and descriptions of the graves

by Béla Miklós Szőke, Sándor Évinger, Csilla Líbor, Frigyes Szücsi, Zsolt Petkes, Balázs Gusztáv Mende, Piroska Rácz, Veronika Csáky

In the following section, we provide a brief overview of the sampled archaeological sites, including detailed descriptions of burials associated with the samples selected for full genome analyses. Information on other samples and negative results is listed in Table S1.

#### 1.1 Zalavár-Vársziget

Zalavár-Vársziget (Zalavár Castle Island, hereafter referred to as Zalavár) was the administrative and ecclesiastical center of the Pannonian Inferior Province in the Carolingian Empire from the 840s CE, and is mentioned in written sources as *Mosaburg, civitas Priwinae*, *urbs paludarum Chezilonis*. In 890 CE, in a grant by King Arnulf, he referred to the Pannonian seat as *regia civitas Mosaburc*. In the Carolingian period, three churches were founded on Castle Island: the first was dedicated to St. John the Baptist in the early 840s CE, the second to the Virgin Mary in 850 CE, and the third to the martyr Hadrian, built between 855-859 CE.

Regular archaeological research on Zalavár-Vársziget (hereafter Zalavár, Fig. S1) began in the early 1950s and has continued uninterrupted since then, with an increase in activity starting in the early 1990s. In the second half of the 19th century and the first half of the 20th century, large parts of the site were quarried for stones and sand, and the church of St. Mary was completely destroyed; therefore its location, size and shape can only be reconstructed on the basis of a 16th century CE drawing. The two cemeteries, consisting of thousands of graves, established around the churches of Mary and Hadrian, situated 50 meters apart, had emerged by the end of the 9th century CE, and the burials of the 10th and 11th centuries CE were dug over the 9th century CE graves, and were partly cut into the earlier graves.

The church of St. John the Baptist was built in the 840s CE for the baptism of a completely pagan local population, therefore no burials were placed around it. The church of St. Mary was a private church in the fortified manor house of the founding Count of Mosaburg, Priwina (or in other sources Pribina) and his son Chezil (also known as Kocel). Around the church only a few people other than his extended family and relatives were allowed to be buried. In contrast, the pilgrimage church of Hadrian the Martyr in the second half of the 9th century CE was surrounded by the graves of noble families and their entourage who were in the immediate vicinity of Priwina and did not have their own mansion and/or private church. This nobility was represented by the wearing of spur sets for men and gold or gilded silver jewelry (earrings, rings) for women, but also by the family burial places surrounded by a stone wall or a wall of planks, the large plank-lined grave pits and the coffins made of heavy planks held together with iron clamps.

From the beginning of the 10th century CE onward, the temples were neglected, and their condition deteriorated. They were surrounded by pagan burials again, and food/drink grave goods (pots, animal bones) appeared in the graves, which were no longer oriented towards the temple. Family ties apparently determined the location of the graves, and it is not coincidental that new groups of graves appear mainly around the pilgrimage temple of Hadrian at this time.

The Western monks who returned in the late 10th and early 11th centuries CE were only able to renovate the church of St. Mary, and when it was rededicated, it was consecrated to St. Adrian. In the 11th century CE, the St. Adrian church of the Benedictine monastery (mentioned as early as 1019 CE) was the burial place of the wealthy families. The jewelry (S-shaped hair rings and rings) of those buried here was made of silver, rarely gold, and silver denarii were often placed in the tombs[5].

**AHP01**
Grave *4/03 - No grave goods; located in the filling of the 'great ditch' (*i.e.* wide, deep fortification ditch protecting the fortified Priwina courtyard (known as *munimen*) from the north). Anthropological and genetic sex: Male; 35-40 years old.

**AHP02**
Grave 49/98 - No grave goods; above settlement object 1/99. Genetic sex: Male; 0,5-1 year old

**AHP03**
Grave 63/95 - No grave goods; Anthropological sex: Male(?); genetic sex: Female; 35-40 years old.

**AHP04**
Grave 77/95 - No grave goods; below feature 55/95, in the filling of the 'great ditch'. Genetic sex: Male; 2-3 years old.

**AHP05**
Grave 78/95/1 - No grave goods; in the filling of the 'great ditch'. Genetic sex: Male; 3-4 years old.

**AHP06**
Grave 79/95 - No grave goods; in the filling of the 'great ditch'. Anthropological and genetic sex: Female; 60-80 years old.

**AHP07**
Grave 93/95 - No grave goods; in the filling of the 'great ditch'. Genetic sex: Male; 4-5 years old.

**AHP08**
Grave 94/95 - No grave goods; in the fill of the 'great ditch'. Genetic sex: Male; 3-4 years old.

**AHP09**
Grave 96/95 - No grave goods; below feature 91/95; in the filling of the 'great ditch'. Anthropological and genetic sex: Male; 25-30 years old.

**AHP10**

Grave 464/91D - Grave good: thin silver wire hoop; above feature 466/91D and 472/91D. Genetic sex: Female; 2-3 years old.

**AHP11**

Grave 466/91E,B - Grave good: iron buckle; below feature 467/91E. Anthropological and genetic sex: Female; 40-80 years old.

**AHP12**

Grave 3/83 - No grave goods; tomb - brick-roofed crypt near the NW corner of Hadrian's Church, c. 860-870, early phase of Carolingian period. Anthropological and genetic sex: Male; 40-50 years old.

**AHP13**

Grave 6/00 - No grave goods; above grave 8/00 - early; also feature 1/2000, dug in the filling of the bell-founding pit, theoretically roughly contemporary with grave 8/00, only the orientation of 6/00 is the same as that of the church. Anthropological and genetic sex: Male; 30-40 years old.

**AHP14**

Grave 24/12 - No grave goods. Anthropological and genetic sex: Male; 30-40 years old.

**AHP15**

Grave 25/00 - The design of the burial pit follows the previous Avar period rite: narrow trenches at the head and foot ends beneath the coffin. The coffin was held together by 10 large iron pins, next to the dead was a small iron knife and a so-called Moravian-type fire-brake (typical in the second half of the 9th century CE, especially in the Moravian Basin). Grave 19/00 cuts into this grave, which contained an ornate gold dangling. This is one of the most precious of the Zalavár jewels, its decoration has parallels on the so-called *Arnulf ciborium* and on the cover of the Codex Aureus, which was made around 870 CE at the court of Charles the Bald in St. Denis and given by Arnulf to the monastery of St. Emmeram in Regensburg in 893 CE. Hence, the individual in grave 19/00 probably belonged to the entourage of King Arnulf, who was a frequent visitor to Mosaburg between 880 and 890 CE. Consequently, grave 25/00 probably is from between 860-870 CE. Anthropological and genetic sex: Male; 40-50 years old.

**AHP16**

Grave 297/03 - No grave goods; dug into settlement object 2/02, coffin grave, Carolingian but possibly early 10th century. Anthropological and genetic sex: Male; 30-35 years old.

**AHP17**

Grave 166/01 - No grave goods; in a larch wood coffin, filling mortared (from putty from church construction), aligned with the wooden platform around the church, early phase of the Carolingian period. Anthropological and genetic sex: Male; 45-50 years old.

**AHP18**

Grave 193A/99 - No grave goods; above 214/99. Carolingian period grave, but its orientation deviates from the average: exactly NW-SE. Anthropological and genetic sex: Male; 50-60 years old.

**AHP19**

Grave 226/99 - No grave goods; below 79/99, 34/99, 225/99. Trench-like depressions at either end of the grave pit, below the coffin. The tomb is in the lowest layer of graves from the early phase of the Carolingian period, with several layers above it, also of Carolingian date and with an Árpádian period burial. Anthropological and genetic sex: Male; 40-50 years old.

**AHP20**

Grave 234/99 - In a coffin with large nails, early Carolingian phase. Anthropological and genetic sex: Male; 40-50 years old.

**AHP21**

Grave 8/00 - Grave goods: two iron knives, stratigraphically below features 119/99, 27/99, 6/00, 28A-B/99, 46/99, from the early phase of the Carolingian period. Divergence of the orientation from SW to NE. Radiocarbon date: 774-992 cal CE (95.4% CI, 1140±30 BP, Fig. S2), based on archaeological observations, the burial of this individual predates the Hungarian conquest. Anthropological and genetic sex: Male; 35-40 years old with a healed left clavicle and a similarly healed left scapula fracture.

This burial belongs to one of the earliest layers of the cemetery but possibly not the oldest. The Hadrianus Church was consecrated in the second half of the 850s, and the grave was dug into the fill of a feature nr. 1/2000 identified as the bell-casting pit of the church. This part of the cemetery is slightly farther from the church and outside the wooden fence, where burials began from the mid-870s when the church's function changed. This grave is also different in its orientation, not aligned with the church. A woman's grave (46/99) with jewelry dating to the last third of the 9th century lies within the fill of the grave pit, suggesting the burial 8/00 around the 880s, but not later than 890.

**AHP22**

Grave 22/01 - No grave goods; below grave 21/01. Probably buried in the last quarter of the 9th century (c. 880-890 CE), as the skeleton was placed above a pillar pit of the bishop’s wooden palace that was demolished c. 875-880 CE. Anthropological and genetic sex: Male; 30-35 years old.

**AHP23**

Grave 51/98 - No grave goods; below of graves 20/98, 21/98, 42/98, 30/98, coffin burial. Trench-like depression in the two narrow ends of the grave below the coffin, a third depression below the pelvic. Early phase of the Carolingian phase, c. 860-870. Anthropological and genetic sex: Male; 45-55 years old.

**AHP24**

Grave 103/99 - No grave goods; early phase of the Carolingian period, immediately below tomb 91/99, also of Carolingian date. Anthropological and genetic sex: Male; 25-35 years old.

**AHP25**
Grave 133/01 - No grave goods; below 134/01. Anthropological and genetic sex: Male; 45-50 years old.

**AHP26**
Grave 169/01 - No grave goods; placed in a carved wooden coffin. Anthropological and genetic sex: Male; 25-30 years old.

**AHP27**
Grave 202/99 – No grave goods; below graves 112/99, 77/99. Early phase of the Carolingian period, ditch-like depressions at both narrow ends of the grave pit. Anthropological and genetic sex: Male; 35-40 years old.

**AHP28**
Grave 225/90 - No grave goods; placed in an ironed coffin below grave 222/90, above graves 219/90 and 226/90. Anthropological and genetic sex: Male; 40-45 years old.

**AHP29**
Grave 232/99 - No grave goods; below graves 228/99, 231/99, 233/91, 176/99, 186/99. Buried in a large coffin with 14 coffin pins and staples, early Carolingian period, c. 860-870. Anthropological and genetic sex: Male; 25-35 years old.

**AHP30**
Grave 277/90 - No grave goods; above the moat of our church, end of Carolingian period, early 10th century CE. Anthropological and genetic sex: Male; 40-50 years old.

**AHP31**
Grave 284/88-90 - Grave goods: spur set. Anthropological and genetic sex: Male; 30-35 years old.

**AHP32**
Grave 343/90-91 + 78/00 - No grave goods; placed in a carved wooden coffin below the graves 342/90, 228/90, and below settlement object 4/91; Anthropological and genetic sex: Male; 30-40 years old.

**AHP33**
Grave 420/91D - No grave goods; below grave 421/91 D. Anthropological and genetic sex: Male; 20-25 years old.

**AHP34**
Grave '7/03 - No grave goods; above grave 76/02, below 30/03. Anthropological and genetic sex: Male; 35-40 years old.

**AHP35**

Grave 9/07 - No grave goods; in grave field, Carolingian glass fragment, next to Árpádian period palisade trench. Anthropological and genetic sex: Male; 40-50 years old.

**AHP36**

Grave 13/08 - No grave goods but animal bone in the grave field found, at the northern edge of the 'big ditch'. Anthropological and genetic sex: Male; 45-55 years old.

**AHP37**

Grave 14/08 - No grave goods; Anthropological and genetic sex: Male; 30-40 years old.

**AHP38**

Grave 25/96 - No grave goods; above the Carolingian palace pillar holes, above grave 26/96, below grave 37/96. Anthropological and genetic sex: Male; 45-55 years old.

**AHP39**

Grave 29/95 - No grave goods; above grave 50/95. Anthropological and genetic sex: Male; 50-60 years old.

**AHP40**

Grave 38/02 - No grave goods; placed in a coffin, early Árpádian period phase. Anthropological and genetic sex: Male; 35-40 years old.

**AHP41**

Grave 74/04 - No grave goods; partial stone dressing. Anthropological and genetic sex: Male; 30-40 years old.

**AHP42**

Grave 79/02 - No grave goods; situated above grave 74/03. Anthropological and genetic sex: Male; 40-45 years old.

**AHP43**

Grave 15/98 - Grave good: bottom-stamped vessel, close to the eastern palisade wall. Anthropological and genetic sex: Male; 45-55 years old.

**AHP44**

Grave 29/98 - Grave good: bronze object, near the base of the palisade wall. Anthropological and genetic sex: Male; 40-50 years old.

**AHP45**

Grave 52/95 - No grave goods; below an Árpádian period pit. Anthropological and genetic sex: Male; 40-50 years old.

**AHP46**

Grave 64/95 - No grave goods; below grave 53/95. Anthropological and genetic sex: Male; 35-40 years old.

**AHP47**
Grave 67/95 - No grave goods. Anthropological and genetic sex: Male; 40-45 years old.

**AHP48**
Grave 69/95 - No grave goods; palisade wall above base ditch. Anthropological and genetic sex: Male; 45-55 years old.

**AHP49**
78/95 Grave - No grave goods. Anthropological and genetic sex: Male; 25-30 years old (?).

**AHP50**
Grave 88/95 - No grave goods; below grave 87/95. Anthropological and genetic sex: Female; 35-40 years old.

**AHP52**
Grave 98/95 - Grave good: iron knife; below grave 72/95. Anthropological and genetic sex: Female; 40-45 years old.

**AHP53**
Grave 124/00 - No grave goods; placed in an ironed coffin, below grave 113/00. Anthropological and genetic sex: Male; 40-50 years old.

**AHP54**
Grave 3/95 - No grave goods; above graves 4/95, 9/95, 5/95. Anthropological and genetic sex: Male; 25-30 years old.

**AHP55**
Grave 6/02 - No grave goods; grave was cut by burial 7/02 (bead row with pierced kauri coils in it), above tombs 5/02, 31/02. Anthropological and genetic sex: Male; 30-35 years old.

**AHP56**
Grave 12/12 - No grave goods; the head of the grave pit is next to the eastern wall of the Arnulf Palace, built in the 880s, the palace was already in ruins when the grave was dug. Anthropological and genetic sex: Male; 35-45 years old.

**AHP57**
Grave 20/96 - No grave goods; below grave 24/96, above grave 85/96, above a narrow brick palisade trench. Anthropological and genetic sex: Male; 45-55 years old.

**AHP58**
Grave 22/96 - No grave goods. Anthropological and genetic sex: Male; 35-40 years old.

**AHP59**

Grave 30/03 - No grave goods; above graves 7/03, 5/03. Anthropological and genetic sex: Male; 25-30 years old.

**AHP60**

Grave 48/98 - No grave goods; sunk into the foundation trench of the slate wall surrounding Hadrian's church. Anthropological and genetic sex: Female; 30-40 years old.

**AHP61**

Grave 51/96 - No grave goods; in the filling of the 'big trench' (*munimen* fortification ditch). Anthropological and genetic sex: Male; 40-50 years old.

**AHP62**

Grave 73/04 - No grave goods; above grave 89/04. Anthropological: Male; genetic sex: Female; 60-80 years old.

**AHP63**

Grave 100/95 - No grave goods. Anthropological and genetic sex: Male; 35-40 years old.

**AHP64**

Grave 197/99 - No grave goods. Anthropological and genetic sex: Male; 25-30 years old.

**AHP65**

Grave 3/08 - No grave goods but animal bone was recovered from grave soil. Anthropological and genetic sex: Male; 30-40 years old.

**AHP66**

Grave 23/12 - No grave goods; above settlement object 3/12 (Carolingian period). Anthropological and genetic sex: Male; 35-50 years old.

**AHP67**

Grave 29/03 - No grave goods. Anthropological and genetic sex: Male; 35-40 years old.

**AHP68**

Grave 74/03 - No grave goods; early Árpádian period, possibly 10th century CE. Anthropological and genetic sex: Male; 45-50 years old.

**AHP69**

Grave 78/01 - No grave goods; stratigraphic evidence suggests Árpádian period dating. Anthropological and genetic sex: Male; 45-50 years old.

**AHP70**

Grave 146/01 - No grave goods. Anthropological and genetic sex: Male; 50-60 years old.

**AHP71**

Grave 285/90 - Grave good: iron nail; below graves 278/88, 279/88, 31/90, two other graves below, also Carolingian, earlier than these. Anthropological and genetic sex: Male; 35-45 years old.

**AHP72**

Grave 466/91E "A" - Grave good: small silver ear-ring with twisted end (10th century CE). Anthropological and genetic sex: Female; 30-50 years old.

#### 1.2 Himod-Káposztásföldek

The site of Himod-Káposztásföldek (hereafter Himod, Fig. S3) is located east of the Amber Road linking *Savaria* (today's city of Szombathely) with *Carnuntum* (Petronell), on the eastern border of *Avaria*, approximately 30 km east of the 9th century CE Sopronkehida cemetery. According to the excavator Péter Tomka, the graves of the Carolingian period in Himod, which were excavated in a narrow, only 7 meters wide strip, were buried in the first two thirds of the 9th century CE (Himod phase I, hereafter Himod I). The site is among the few early medieval commoner cemeteries in the Carpathian Basin where the graves of the 10th-11th centuries CE (Himod II) were superposed on the 9th century CE ones[50], [51].

**KOZ1**

Grave 104 - Grave goods: beads; excavated on settlement object 31, above grave 103. Genetic sex: Female; 9-12 years old.

**KOZ2**

Grave 117 - Grave good: pot fragment; grave 129 was layered on it. Genetic sex: Female; 1-6 years old.

**KOZ3**

Grave 127 - Grave goods: pot, small bronze ring. Genetic sex: Female; 7-9 years old.

**KOZ4**

Grave 129 - Grave good: pot with bottom stamp; above grave 117. Genetic sex: Female; 8-12 years old.

**KOZ5**

Grave 153 - Grave goods: hair ring, bronze ring; above grave 152. Genetic sex: Female; 10-14 years old.

**KOZ6**

Grave 155 - Grave goods: silver earrings. Genetic sex: Female; 1.5-3 years old.

**KOZ7**

Grave 140 - Grave goods: silver band, ring, pearl, iron knife, disc earrings; below grave 139. Anthropological and genetic sex: Female; 25-29 years old.

**KOZ8**

Grave 144 - Grave good: bronze ring. Anthropological sex: Male? (fragmented skeleton); genetic sex: Female; 40-59 years old.

**KOZ9**
Grave 146 - Grave goods: light stricker iron, iron knife, pot. Anthropological and genetic sex: Male; 35-44 years old.

**KOZ10**
Grave 151 - No grave goods. Genetic sex: Female; 0.5-1.5 years old.

**KOZ11**
Grave 100 - No grave goods; above prehistoric pit. Anthropological and genetic sex: Female; 30-39 years old.

**KOZ12**
Grave 128 - Grave goods: bronze strap ring, iron knife. Anthropological and genetic sex: Female; 30-39 years old.

**KOZ13**
Grave 136/B skeleton - Grave good: iron knife; below the grave 135/A. Anthropological and genetic sex: Female; 40-49 years old.

**KOZ14**
Grave 142 - Grave goods: spearhead, gilded silver buckle, iron buckle, iron belt end, iron knife, arrowheads, iron hoop, other iron objects (fire making tools), pot, "*Wadenbindengarnitur*". Anthropological and genetic sex: Male; 40-49 years old.

**KOZ15**
Grave 42 - Grave goods: S-end hoop jewelry (ear- or hair-ring). Genetic sex: Female; 1.5-3 years old.

**KOZ16**
Grave 57 - Grave goods: knife, arrowheads. Anthropological and genetic sex: Male; 20-39 years old.

**KOZ17**
Grave 78 - Grave goods: twisted-end hoop jewelry, bronze needle. Genetic sex: Male; 5-7 years old.

**KOZ18**
Grave 83 - Grave goods: bow handle bone brace, silver ball button, arrowhead, animal bone. Anthropological and genetic sex: Male; 50-59 years old.

**KOZ19**

Grave 93 - Grave goods: two-piece dangling, ring, beads, hoop jewelry. Genetic sex: Female; 15-19 years old.

**KOZ20**
Grave 82/A - Grave good: plain open bracelet. Anthropological and genetic sex: Female; 50-59 years old.

**KOZ21**
Grave 84 - Grave goods: bracelet, bangle, ring. Anthropological and genetic sex: Female; 40-44 years old.

**KOZ22**
Grave 105 - Grave good: silver band ring. Genetic sex: Female; 8-12 years old.

**KOZ23**
Grave 109 - Grave good: hoop jewelry with twisted ends. Anthropological and genetic sex: Female; 40-49 years old.

**KOZ24**
Grave 113 - Grave goods: silver and bronze hoop jewelry. Anthropological and genetic sex: Female; 25-34 years old.

**KOZ25**
Grave 115 - Grave goods: earring fragments - bronze hoop, hoop jewelry. Anthropological sex: Female; genetic sex: Male; 20-24 years old.

**KOZ26**
Grave 119 - Grave goods: arrowhead, hoop jewelry, knife. Anthropological and genetic sex: Male; 25-34 years old.

**KOZ27**
Grave 138 - Grave goods: iron object, bronze hoop, thick S-ended hairband; the grave pit is contiguous with grave 149. Anthropological and genetic sex: Female; 40-59 years old.

**KOZ28**
Grave 116 - Grave good: iron knife. Anthropological and genetic sex: Male; 40-49 years old.

#### 1.3 Sárbogárd-Tringer tanya

This site was discovered during the construction of a road in early March 1961. During the subsequent excavation, between 16 March and 7 July 1961, a medieval cemetery composed of a total of 100 graves was uncovered (Fig. S4). The cemetery may have been used only in the 10th century CE. In the early-dated graves, typical Conquest period burial practices (symbolic horse burials) and artifacts (weapons: bow, arrowhead; female jewelry: hair braiding discs, earrings with ball-shaped dangles) were observed. In contrast, toward the end of the century, the graves became much poorer, containing only a few wire hoops and crescent-shaped danglers, artifacts which became widespread in the last third of the 10th century CE. There were also significant differences in the orientation of the graves and their positioning in relation to each other between the earlier and later phases. While the early-dated graves were mostly oriented northwest to southeast and somewhat irregularly spaced, the late-phase graves were oriented southwest to northeast and arranged in approximately regular rows. The early-phase graves were located in the northern part of the cemetery, while the late graves were in the southern part. However, a few late-phase burials appeared in the northern part of the cemetery, wedged between the early graves without disturbing them[52], [53], [54].

**AHP117**
Grave 5 - Symbolic horse burial, side-plate oats for a foal, six plates of bone stiffening of a bow, bone plate of bow stiffener, bow grip plate, round iron buckle, pear-shaped iron stirrup, two iron stiffening ears of a quiver, arrowhead. Anthropological and genetic sex: Male; 47-51 years old.

**AHP118**
Grave 6 - No grave goods. Anthropological and genetic sex: Male; 51-55 years old.

**AHP119**
Grave 7 - No grave goods. Anthropological and genetic sex: Male; 35-39 years old.

**AHP120**
Grave 12 - Grave good: iron knife. Anthropological and genetic sex: Female; 47-53 years old.

**AHP121**
Grave 17 - No grave goods. Anthropological and genetic sex: Male; 19-22 years old.

**AHP122**
Grave 19 - Grave goods: bronze strap bracelet with twisted end, iron fragment. Anthropological and genetic sex: Female; 17-21 years old.

**AHP123**
Grave 22 - No grave good. Anthropological and genetic sex: Male; 41-47 years old.

**AHP124**
Grave 24 - Grave goods: spherical dangle earrings, bronze chain link with small bronze plate, silver plated bronze hair braid discs with centrally arranged palmette decoration, kauri snails and beads of various colors, types and materials, bronze plate ball button, bronze band ring, silver mounting. Genetic sex: Female; ~14 years old.

**AHP125**
Grave 25 - No grave goods. Anthropological and genetic sex: Female; 55-61 years old.

**AHP126**
Grave 26 - No grave good. Anthropological and genetic sex: Male; 56-60 years old.

**AHP127**
Grave 30 - Grave goods: bronze hoop with beads, lance-shaped iron pliers, iron knives, flint, pith. Anthropological and genetic sex: Male; 30-36 years old.

**AHP128**
Grave 40 - No grave good. Anthropological and genetic sex: Female; 54-60 years old.

**AHP129**
Grave 41 - Grave good: iron knife. Anthropological and genetic sex: Male; 19-22 years old.

**AHP130**
Grave 46 - Grave goods: bronze hoop, bronze bracelet, bronze ring, piece of cattle or horse pelvic bone; grave placed into the western end of grave 52. Anthropological and genetic sex: Female; 60-70 years old.

**AHP131**
Grave 48 - No grave goods. Anthropological and genetic sex: Male; 23-30 years old.

**AHP132**
Grave 49 – Grave good: dress decoration made of pressed bronze plate. Genetic sex: Female; 4.5-5 years old.

**AHP133**
Grave 53 - No grave goods. Anthropological and genetic sex: Male; 20-22 years old.

**AHP134**
Grave 55 - Grave goods: iron buckle, iron knife. Anthropological and genetic sex: Male; 36-40 years old.

**AHP135**
Grave 57 - Grave goods: eggshell, bronze strap ring, bronze wire ring. Genetic sex: Male; 5-5.5 years old.

**AHP137**
Grave 64 - Grave goods: iron shards, iron bracelet. Anthropological and genetic sex: Male; 38-42 years old.

**AHP138**
Grave 66 - No grave goods. Anthropological and genetic sex: Male; 50-56 years old.

**AHP139**
Grave 72 - No grave goods. Anthropological and genetic sex: Male; 48-52 years old.

**AHP140**

Grave 80 - Grave good: bronze wire ring. Anthropological and genetic sex: Female; 55-59 years old.

**AHP141**
Grave 9 - Grave goods: S-ended silver rings, bronze wire ring. Anthropological and genetic sex: Female; 62-70 years old.

**AHP142**
Grave 10 - No grave goods. Anthropological and genetic sex: Female; 62-75 years old.

**AHP143**
Grave 15 - No grave goods. Genetic sex: Male; ~6 years old.

**AHP144**
Grave 23 - No grave goods. Genetic sex: Male; ~3 years old.

**AHP145**
Grave 31 - Grave goods: cylindrical mineral beads. Genetic sex: Male; 1.5-2 years old.

**AHP146**
Grave 32 - Grave good: rectangular iron buckle. Anthropological and genetic sex: Female; 65-75 years old.

**AHP148**
Grave 38 - No grave goods. Anthropological and genetic sex: Male; 65-75 years old.

**AHP149**
Grave 50 - Grave good: arrowhead. Anthropological and genetic sex: Male; 52-56 years old.

**AHP150**
Grave 68 – Grave goods: S-end silver rings, bronze wire ring. Anthropological and genetic sex: Female; 55-61 years old.

**AHP151**
Grave 71 - No grave goods. Anthropological and genetic sex: Male; 54-58 years old.

**AHP152**
Grave 74 - No grave goods. Anthropological and genetic sex: Male; 32-36 years old.

**AHP153**
Grave 75 - No grave goods. Anthropological and genetic sex: Male; 44-48 years old.

**AHP154**
Grave 78 - No grave goods. Anthropological and genetic sex: Male 17-18 years old.

**AHP155**
Grave 83 - Grave goods: S-end bronze rings. Anthropological and genetic sex: Female; 46-52 years old.

**AHP156**
Grave 84 - Grave goods: bronze crescent shaped dangling, green mineral bead. Genetic sex: Female; 9-9.5 years old.

**AHP158**
Grave 87 - Grave goods: beads. Genetic sex: Female; ~5.5 years old.

**AHP159**
Grave 90 - Grave goods: S-shaped bronze rings. Anthropological and genetic sex: Female; 51-57 years old.

#### 1.4 Visegrád

##### 1.4.1 Visegrád-Széchenyi street 25.

The site is a part of a commoner cemetery (land parcel No. 322) dated to the mid-11th century CE, excavated in 2008 by István Kováts and Péter Gróf (Hungarian National Museum - Matthias King Museum/Mátyás Király Museum) on the left bank of the Apátkúti Creek in Visegrád. The burials were placed in a row: a total of five early Árpádian period graves (Grave 2008/1 - Grave 2008/4, Grave 2008/7) oriented west to east, and at least three other graves were found as isolated finds[55].

**AHP73**
Grave 2008/1 – Grave goods: bronze hair ring near left temple, bronze patina remains on some finger bones. Anthropological and genetic sex: Female; Adultus age.

**AHP74**
Grave 2008/2 - No grave goods. Anthropological and genetic sex: Male; Adultus-maturus age.

**AHP75**
Grave 2008/3 - Grave goods: two bronze hoop jewels and cylindrical and lenticular glass beads at right shoulder and on the chest. Anthropological and genetic sex: Female; Adultus age.

**AHP76**
Grave 2008/4 - Grave goods: bronze hoop rings on both ring fingers. Anthropological and genetic sex: Female; Adultus age.

**AHP77**
Grave 2008/7 - Grave goods: left side of head a bronze S-ended hoop jewel and a plain ring was placed. Genetic sex: Female; Infans II age (late childhood).

##### 1.4.2 Visegrad-Sibrik Hill, site I

An early medieval cemetery (or cemetery complex) was excavated to the southeast of the late Roman fortress on Sibrik Hill. The cemetery(s) and the associated church(es) (or a two-phase church) were excavated by Mátyás Szőke in 1972-1974 (graves) and 1977-1979 (graves and a building complex). It is not clear whether the 221 recovered graves belong to one or two cemeteries: a part of a row (?) cemetery (146 graves) is recognisable, and to the east of it a two-phase church (church 1: dated to the foundation of the state, *i.e.* contemporaneous with the construction of the *ispanic*[1] center, church 2: from the mid-11th to the mid-12th centuries CE) and its associated burials (75 graves). Whether the 11th century CE cemetery, characterized by graves arranged in a row, is connected to the cemetery surrounding the second church, thereby forming a single burial ground, remains an open question. In both the row-type cemetery and on the western side of church 2, groups of graves have been identified, which, according to the artifacts found, appear to have belonged to important members of the community. Coins (Solomon, Géza *dux*, Géza I, László I), 'hair rings' (hoops), pearl necklaces, arrowheads in the row-type cemetery, a medal (László I) and gold and silver 'hair rings' and rings in the section around the church, and a Roman-period column fragment placed as a grave marker to one of the graves indicate the higher social status of the people buried there[56]. The gold rings from the graves of two men and a woman near the church are paralleled with the gold ring in graves of Sibrik Hill II cemetery (see individual AHP114, grave 2013/1). The dateable coins (Solomon, Géza I, László I) suggest that both groups of graves were established at approximately the same time, but they differ slightly in their burial customs (in terms of grave goods). The rich graves next to the second church could be the burial place of the noblemen of the castle, or even the *ispan* himself and his family. A third group of well-furnished clergymen graves, those who served the temple, were found behind the sanctuary of church 2. Three relocated (exhumed) graves have been documented next to the church, which are presumably related to the exhumed masonry tomb with a gold ring from Sibrik Hill II[57].

From an art historical point of view, the small columns of the gallery place the building in contemporaneity with the nearby St. Andrew's Monastery, and it is easy to imagine that the same masters worked on the design of both[57]. Based on the fragments of frescoes that have been found, many assume Byzantine connections[58], [59].

**AHP78**

Grave 76/137 (uncertain phase) - Grave goods: 3 fragments of knives and a sword. Anthropological and genetic sex: Female; Maturus age.

**AHP79**

Grave 77/167 (phase 0/I?) - No grave goods. Anthropological sex: Female; Genetic sex: Male; Maturus age.

**AHP81**

---

[1] Ispan or in Lat. *comes* was an official who governed a county (comitatus) on behalf of the king in historical Hungary.

Grave 79/214 (phase 0/I) - No grave goods; superposition: located under the floor of the church 2. Filling yellowish brown soil mixed with charcoal in some places. Anthropological sex: Female; Genetic sex: Male; Juvenis age.

**AHP82**
Grave 79/215 (phase 0/I) - No grave goods; superposition: located under the floor of church 2. Filling is yellowish brown earth mixed with charcoal in some places. Genetic sex: Female; Infans II age (late childhood).

**AHP84**
Grave 79/221 (phase 0/I) - No grave goods; superposition: located under the floor of church 2. Filling is yellowish brown soil mixed with charcoal in some places. Anthropological and genetic sex: Male; Maturus age.

**AHP85**
Grave 74/92 (phase II) - Grave goods: silver coin of Prince Géza under the left hand (CNH.I.23). Anthropological and genetic sex: Female; Maturus age.

**AHP86**
Grave 74/93 (phase II) - Grave goods: silver coin of St. Ladislas I under the right hand (CNH.I31), 2 s-shaped hair rings; superposition: intercut grave 74/120. Anthropological and genetic sex: Male; Maturus age.

**AHP87**
Grave 74/109 (phase II) - Grave goods: coin of Prince Géza (CNH.I23), hair rings. Anthropological sex: Male; Genetic sex: Female; Maturus age..

**AHP91**
Grave 77/164 (phase II) - Grave goods: chalice, paten. Anthropological and genetic sex: Male; Maturus age.

**AHP92**
Grave 77/181 (phase II) - No grave goods. Anthropological and genetic sex: Male; Maturus age.

**AHP93**
Grave 77/186 (phase II) - Grave goods: coin under the left hand in the pelvic. Anthropological and genetic sex: Female; Maturus age.

**AHP94**
Grave 77/188 (phase II) - Grave good: S-shaped hair ring. Anthropological and genetic sex: Female; Juvenis age.

**AHP96**
Grave 74/7 (phase II) - Grave goods: S-shaped hair rings: silver on the left side, tin on the right side. Genetic sex: Female; Maturus age.

**AHP97**
Grave 74/58 (phase II) - Grave good: knife with steel edge, above the left leg. Genetic sex: Male; Infans II age (late childhood).

**AHP98**
Grave 74/110 (phase II) - Grave goods: coin of St. Ladislas I (CNH.I.31), 2 S-shaped hair rings, iron nail. Anthropological and genetic sex: Female; Adultus age.

**AHP99**
Grave 77/153 (phase II) - Grave goods: silver S-shaped hair ring. Genetic sex: Female; Infans II age (late childhood).

**AHP100**
Grave 79/216 (phase 0/I) - No grave goods; superposition: located under the floor of the church II. Filling is yellowish brown earth mixed with charcoal in some places. Anthropological and genetic sex: Female; Maturus age.

**AHP101**
Grave 79/218 (phase 0/I) - No grave goods; superposition: located under the floor of Church II. Filling yellowish brown earth mixed with charcoal in some places. Anthropological and genetic sex: Female; Juvenis age.

**AHP103**
Grave 77/174 (phase 0/I) - No grave goods. Genetic sex: Male; Infans I age.

**AHP104**
Grave 77/175 (phase 0/I) - No grave goods. Anthropological and genetic sex: Female; Juvenis age.

**AHP107**
Grave 77/209 (phase 0/I) - No grave goods. Anthropological and genetic sex: Male; Maturus age.

**AHP108**
Grave 77/202 (uncertain phase) - No grave goods. Anthropological sex: Female; Genetic sex: Male; Juvenis age.

**AHP109**
Grave 74/91 (uncertain phase) - Grave goods: hair ring, beads, animal tooth next to the coffin. Anthropological and genetic sex: Female; Juvenis age.

**AHP110**
Grave 72/52 (uncertain phase) - No grave goods. Anthropological and genetic sex: Male, Maturus age.

**AHP111**
Grave 74/114 (uncertain phase) - No grave goods. Anthropological and genetic sex: Male; Adultus age.

##### 1.4.3 Visegrád-Sibrik Hill, site II

The three graves from the foundation period, excavated in the inner area of the Sibrik fortress, can be interpreted in the context of the fortress chapel. The burials, which date back (based on $^{14}C$) to the beginning of the 11th century CE, were excavated in 2013 (excavation leaders: Gergely Buzás, Katalin Boruzs and Szabina Merva): one grave from the interior of the church, presumably the grave of the founder, one grave north of the church with grave goods indicating a priest's grave, and a third grave east of the church without burial finds[57].

**AHP115**
Grave 2013/2/2: late-10th century/early-11th century (DeA-3955, I/825/3b, 1078±25, cal CE 900-1020 (2 sigma level CI)) - Grave good: the left hand held a pewter chalice, around the right hand a 10 cm wide patch of corroded tin fragments are likely the remains of a paten. Anthropological and genetic sex: Male; Adultus-Maturus age.

**AHP116**
Grave 2015/17/1 - No grave goods. Anthropological and genetic sex: Male; Adultus age.

#### 1.5 Sites of Székesfehérvár area

##### 1.5.1 Székesfehérvár-Rádiótelep

The Rádiótelep cemetery (or cemeteries) was located on a peninsular hilltop surrounded by marshland on the southwestern border of Székesfehérvár and was in use from the second half of the 10th to the beginning of the 11th centuries CE.

In 1923, on the northern side of the radio station (Rádiótelep site), the grave of a horseman (with a sword and an axe with handle) was discovered during digging work. Following a few new burials, an excavation in the following year yielded another 61 graves, bringing the total number of burials on the northern side of the hilltop to 66. According to the excavator, the extent of the cemetery was delimited by the rim of the hill that emerged from the marshland[60], [61].

**AHP161**
Grave 43 - Grave goods: remains of a wooden bucket with iron ears, braced with iron bands, rhombus-shaped shirt decoration beads edged with small white beads, cylindrical bronze dangling, ribbed bead, pyramid-shaped glass beads with gold setting at the bottom, two half-eared knobs with hollowed design, leather remains with boot covers. Anthropological and genetic sex: Female; 50-60 years old.

**AHP163**
Grave 7 - Grave goods: oval shaped, circular in section, with pointed end, open bronze wire bracelet. Anthropological and genetic sex: Female; 35-40 years old.

**AHP164**

Grave 9 - Grave goods: oval open bronze hoop, iron knife. Anthropological sex: Male(?); Genetic sex: Female; age unknown.

**AHP169**

Grave 23 - Grave goods: iron knife, circular open bronze hoop. Anthropological and genetic sex: Male; 40-45 years old.

**AHP173**

Grave 33 - Grave goods: three oval open bronze hoops. Anthropological and genetic sex: Female; 25-30 years old.

**AHP174**

Grave 39/a - No grave goods. Anthropological and genetic sex: Female; 30-35 years old.

##### 1.5.2 Székesfehérvár-Sóstó, -Homokbánya, -Vízművek and -Sárkeresztúri út

Different parts of the same cemetery were excavated at different times and published under different names. It was situated on the eastern bank of Sóstó, on a sand hill rising towards the Sóstó creek, in the immediate vicinity of a bridge that was already present in the Middle Ages. The cemetery was used from the second half of the 10th to the beginning of the 11th centuries CE.

Székesfehérvár, Sárkeresztúri út

At the beginning of 1916, soldiers on the southern side of the "Calvinist cemetery" on Sárkeresztúri út found a colt's bridle, two stirrups and a buckle ring (grave 1), in addition to human and horse bones during trench digging. Subsequently, in the spring of 1916, Arnold Marosi discovered 30 more early Árpádian period burials. Of the excavated graves, 13 contained grave goods[62].

Székesfehérvár - Sóstó, Homokbánya

In 1956, a grave from the Conquest period was discovered during sand mining on the western side of the road to Sárbogárd. During the subsequent excavation, Jenő Fitz authenticated the location of the grave, but the anthropological material and other finds were not recovered. Two more partially disturbed graves were found nearby [63].

Székesfehérvár - Sóstó, Vízművek

In 1971, Alán Kralovánszky carried out an excavation and recovered another 12 (or 13) burials, 10 of which were severely disturbed[64].

**AHP177**

Székesfehérvár-Sóstó, Homokbánya Grave 2 - Grave goods: iron pin, chisel, flint, iron knife, iron hoops, rhombic arrowheads. Anthropological and genetic sex: Male.

**AHP183**

Székesfehérvár-Sárkeresztúri út Grave 1 - Grave goods: single-edged, straight iron knife. Genetic sex: Male.

**AHP184**
Székesfehérvár-Sárkeresztúri út Grave 2 - Grave goods: trapezoidal shouldered stirrups, large-hooped chicory with flat oatcakes, iron buckle. Genetic sex: Male.

**AHP185**
Székesfehérvár-Sárkeresztúri út, Grave 3 - Grave goods: open bronze strap rings. Genetic sex: Male.

**AHP187**
Székesfehérvár-Sárkeresztúri út, without grave number (the inventory number: 70.97.10). Anthropological and genetic sex: Male.

##### 1.5.3 Székesfehérvár-Táci út repülőtér (airport)

In 1944, during the construction of the Táci út airport, three burials dating back to the Conquest Period (10th-11th centuries CE) of Hungary were discovered[60].

The graves were presumably not documented due to the wartime situation, so not only is the exact location of the cemetery unknown, but also the circumstances of their discovery and placement in the museum are unclear. It can be assumed that Árpád Dormuth carried out excavations or observations in the area. All of the graves were disturbed, and only some of the finds from the graves have been recovered and delivered to the museum.

**AHP181**
Grave 2 - Grave good: trapezoidal stirrup. Anthropological and genetic sex: Male; 34-38 years old.

##### 1.5.4 Székesfehérvár-Sóstó (Ikarus factory)

Museum specialists led by Alán Kralovánszky were able to excavate four graves dating to the 8th century CE oriented NW-SE from the Avar period cemetery, which had been disturbed during factory construction in 1968. They found dangling bronze earrings, an iron buckle, a pot and animal bones as grave findings[65], [66].

**AHP235**
Grave 2 - Grave goods: vessel, chicken bone, cattle bone, iron buckle, dangling bronze earrings. Genetic sex: Female.

#### 1.6 Bodajk

##### 1.6.1 Bodajk-Proletárföldek

In 2016, a trial excavation revealed 14 10th century CE and 14 Celtic graves on the western side of the Móri Creek, about 250-300 meters from the stream. The size and extent of the

10th century CE cemetery are unknown and could not be delimited in either direction during the trial excavation. Due to the small size of the excavation, the relationship between the graves and the layout of the cemetery cannot be revealed, but the graves were quite irregularly spaced in the excavated area. A satellite image of the area likely shows additional grave patches, and as such the excavated graves could be part of a larger cemetery. The Homoki-dűlő Avar cemetery lies 300-350 meters southeast of the cemetery [67].

**AHP190**
Grave 2 - No grave goods; Genetic sex: Male; 14-16 years old

**AHP194**
Grave 6 - No grave goods; Anthropological and genetic sex: Male; 30-34 years old

**AHP195**
Grave 7 - Grave good: iron knife; the grave was dug over the north-eastern side of an earlier Celtic cremation grave (feature 12), it was robbed. Anthropological sex: Female (?); Genetic sex: Male; 30-80 years old.

**AHP196**
Grave 8 - Grave good: iron knife; dark discoloration of soil next to the body, presumably the mark of a coffin. Disturbance in the grave of uncertain origin concerning the skull and its immediate surroundings. Anthropological and genetic sex: Male; 30-34 years old.

**AHP199**
Grave 18 - Grave goods: bronze strap rings. Genetic sex: Male; 3-4 years old.

**AHP200**
Grave 12 - No grave goods. Genetic sex: Male; 11-12 years old.

**AHP201**
Grave 13 - Grave good: Iron knife. The grave was dug over the northern corner of an earlier Celtic skeletal grave (feature 24). Anthropological and genetic sex: Male; 15-18 years old.

**AHP202**
Grave 14 - No grave goods, disturbed; the grave was dug over two earlier Celtic cremation graves: the western side of Celtic cremation grave SE 22 and the eastern side of Celtic cremation grave SE 28. Anthropological and genetic sex: Male; 30-34 years old.

##### 1.6.2 Bodajk-Homoki dűlő

Of the four known Avar period burial sites around the town of Bodajk, the Homoki-dűlő site, located north of the town, can be located in a satellite image as a cemetery of at least 1,000 graves. The cemetery was in use from the last third of the 6th to the 9th centuries CE, throughout the Avar period, according to the current findings and further stratigraphic observations.

In 2010 and 2011, the staff of the Szent István Király Museum carried out rescue excavations in the area and in 2016, together with the Early Hungarians Research Team of the RCH, Hungarian Academy of Sciences, they continued the excavation of the Avar period cemetery and the 10th century CE and La Tène cemeteries located 200-300 meters away. Over the three excavation seasons, 154 graves were uncovered in the Avar period cemetery, initially under the leadership of Péter Prander and Eszter Pásztor, and later Frigyes Szücsi.

Of the 41 men buried, around every fifth had a decorated mounted belt. Nine graves contained belt sets with plate and pressed bronze or silver ornaments or chased iron and cast bronze belt ornaments, and two belt sets made of rare antler carvings. Weapon accessories are scarce: only one spearhead, five arrowheads in two graves and the antler stiffening plates of a bow were recovered. The seven iron knives (>18 cm blade length) and an iron ax can be interpreted as both tools and weapons. Earrings were found in 52 female graves, beads in 26 graves, bracelets in 3 graves and rings in 2 graves. Twelve graves also contained wheel-thrown pottery, half of them belonged to the so-called 'Csákberény' group.

In terms of Germanic cultural elements (object types, decorative elements), the cemetery at Homoki-dűlő can be compared with the unpublished cemeteries at Mezőfalva-vasútállomás and Tác-Fövenypuszta, together with the large East Transdanubian Merovingian type cemeteries (Budakalász, Szekszárd-Bogyiszlói út, Kölked-Feketekapu, Zamárdi), and the nearby Csákberény- Orondpusztai cemetery[68].

**AHP203**
Grave 2 - Grave goods: bronze earrings with overlay decoration, iron buckle, animal bones; dated to the 8th-9th centuries CE. Anthropological and genetic sex: Female; 50+ years old.

**AHP204**
Grave 4 - Grave goods: trapezoid and rectangular iron buckles, animal bone; dated to the 8th-9th centuries CE. Anthropological and genetic sex: Male; 50-54 years old.

**AHP206**
Grave 19 - Grave goods: amorphous iron object, bronze strap ends, rectangular iron buckle, iron knife, bone comb, iron hoop, animal bone, boat-shaped iron plate, bronze plates, bronze hoop; dated to the middle third of the 7th century CE. Anthropological and genetic sex: Male; 30-39 years old.

**AHP207**
Grave 23 - No grave goods; dated to the 8th-9th centuries CE. Anthropological and genetic sex: Male; 40-44 years old.

**AHP208**
Grave 40 - Grave goods: Pair of earrings with disc dangles; dated to the 8th-9th centuries CE. Anthropological sex: Male; 40+ years old. Genetic sex: Female.

**AHP209**

Grave 44 - Grave goods: iron buckle, iron rings, iron knife, animal bone; dated to the 8th-9th centuries CE. Anthropological and genetic sex: Male; 25-29 years old.

**AHP211**
Grave 117 - Grave goods: earring, bead, iron buckle, iron hoop, animal bone; last third of the 7th century CE. Anthropological and genetic sex: Female; 40-44 years old.

#### 1.7 Előszállás-Bajcsihegy (Mezőfalva-Vasútállomás)

The first findings were recovered by the MÁV railway company in 1931 on the northern slope of Bajcsihegy. Unfortunately, most of the graves were lost, but Arnold Marosi estimated the number of destroyed graves at 30-40. In the same year, eight Avar graves were excavated on the eastern side of the Bajcsihegy hill, and four on the western side[69]. During construction in 1952, the new railway cut through Bajcsihegy and hundreds of graves were destroyed, only five were initially recovered, making it necessary to excavate the endangered area. Ultimately 252 graves were uncovered between 1952-1953. Jenő Fitz estimated the number of graves to be at least 2,000, of which about half had been destroyed by the railway construction works[70].

Excavations were carried out in connection with the construction works: 1931 - Nándor Fettich, graves A-C; 1933 - L. Apor, graves 1-12; 1952 - Béla Gergely, graves 13-17; 1952 - Jenő Fitz, graves 18-109; 1952 - István Bóna, graves 110-138; 1953 - Jenő Fitz, graves 139-179; 1953 - István Bóna, graves 180-270. So far, a total of 273 graves have been excavated and 23 vessels have been recorded as isolated finds among the destroyed graves. There were 16 horse and horseman graves among the burials, six of which were in a row and richly equipped with silver fittings[71]. A considerable amount of pottery was found as grave goods, 23 of which were wheel-thrown gray pottery[72]. The findings suggest that the cemetery was used throughout the Avar period[71], [73]. The anthropological material was processed by Sándor Wenger[74].

**AHP220**
Grave 51/A - Grave goods: bronze chain, iron knife, iron buckles; dated to the 7th century CE. Anthropological and genetic sex: Male; Adultus age.

**AHP221**
Grave 109 - Grave goods: animal bones, bronze harness, bronze buckle, black small pot, red-eared pot, iron horn, bronze phalera, bronze plate, rosette horse harness, strap bander, iron buckle, strap end, iron stirrup, strap buckle, strap adjuster, hanging strap, iron arrowhead, iron part for saddle rack, rosette harness for farm ring. The horse grave (man and horse in a common grave pit) is dated to the 7th century CE. Anthropological and genetic sex: Male; Adultus age.

#### 1.8 Csákberény

##### 1.8.1 Csákberény-Orondpuszta

The site was discovered in 1935 during the construction of a railway track. Between 1936-1939 a total of 452 graves were excavated, initially under the direction of József

Lencsés, and later under the supervision of Gyula László. Of the burials excavated, 14 were graves with a horse, 3 were symbolic horse graves (horse implements next to the human body) and 3 contained only a horse. The cemetery was created by groups with different cultural roots in the 6th and 7th centuries CE. Horse graves, weapons of eastern origin, trappings, and harnesses, represent immigrants from the eastern steppes. The Late Antique fibulae placed in the wearing position, the spindle hoops placed on the chest, the bracelets with snake heads and a large number of iron bracelets, as well as the high-quality ceramic vessels with a characteristic red shade (so-called Csákberény group ceramics) made on a high-speed wheel dating back to late Roman workshop traditions (the so-called Csákberény group ceramics), as well as the generally large quantities of ceramic vessels with a quick wheel, suggests the presence of a group with a Romanised culture alongside the Avars. Finally, the number of artifacts of Merovingian-Germanic origin found in the cemetery demonstrates the community's close contact with the Germanic peoples of Western Europe and Northern Italy, raising the possibility of the presence of Germanic communities[75].

In 2019, a total of 53 (453-505) Middle Avar period graves (second half of the 7th - early 8th centuries CE) were excavated during an archaeological survey led by Frigyes Szücsi. Noteworthy finds include bronze strap-ends with pointed ends, various types of belt-plates, earrings with cylindrical-shaped pendulum, and beads, a few melon-seed-form beads and flattened spherical beads with bumpy overlay, necklace ornaments made of lead (including a cross), bronze bells, antlers, and a lash handle end carved from bone. Tomb-like structures were observed in 33 of the 53 graves[76]. An additional 35 graves (506-540) belonging to the 7th century CE were excavated during the 2020 survey. In the latter, a complete set of leg coils (2 bronze large strap ends, 2 bronze small strap ends, 2 bronze buckles) and a nearly intact pressed bronze disc fibula, between the knees, were found. This time only so-called pile-structure graves (2-2 or 3-3 piles found in the longitudinal sides of the grave) were observed, which occurred in a distinct group in the SW part of the section[77].

**AHP229**

Grave 350 - Grave goods: iron knives, iron hoop, iron chisel, a fire-stone, a piece of iron, and a pebbled-lined, thick-walled, ovoid-bodied, narrow mouth, flared, flanged rim, ring handles, unturned, reddish-gray pot and a small pot with a jagged rim, is blackish-yellowish-gray color; dated to the last third of the 6th - first half of the 7th centuries CE. Anthropological sex: Female; Genetic sex: Male; 53-59 years old.

**AHP231**

Grave 390 - Grave good: bronze bracelet; dated to the second third of the 7th century CE. Anthropological and genetic sex: Female; 35-41 years old.

##### 1.8.2 Csákberény-Arató szérű

Between 2021 and 2022, an excavation led by Frigyes Szücsi yielded a total of 85 graves, where two were double (10 and 39) and one was a triple burial (45). The orientation of the graves varied between W-E and NW-SE, with only one skeleton being in the opposite orientation (grave 10). At the feet of individuals, there is pottery. In addition, small, beaded, granulated, tiny, spherical and spiral tipped late earrings, as well as simple earrings, and various late glass beads typically in the shape of millet and/or melon seeds, were common in the graves. Rings, bracelets and animal bones, which can be interpreted as food, were also recovered. Weapons, such as bows, arrowheads and L-shaped axes were uncommon. The cemetery was used between the last third of the 7th to the first half of the 9th centuries CE, and contained no typical Early Avar period artifacts.

**AHP256**

Grave 31 - Grave goods: bronze spiral earrings, iron rings, iron knife, glass beads; dated to the first half of the 9th century CE. The grave is located on the NW edge of the cemetery, adjacent to grave 27 with a late Avar-period cast bronze lily-decorated belt set. Genetic sex: Female; child.

#### 1.9 Pákozd-Börgöndpuszta (Székesfehérvár-Börgöndpuszta)

The site is located on the edge of the marshland of Lake Velence on a hillside facing eastward. In 1960, Alán Kralovánszky excavated graves in two phases. The grave finds consist of vessels, including yellow ceramics, cast bronze mounts with gryphon decorations, iron knives, as well as bronze and iron buckles. All of the excavated graves were undisturbed[78].

**AHP245**

Grave 9 - Grave goods: medium silted pot with notches on the rim, irregular wavy decoration on the upper part, gray color, sooty outside, bone with bronze rivets, animal bones; dated to the 8th century CE. Anthropological and genetic sex: Female; Adult age

**AHP249**

Grave 19 – Grave goods: hen's egg, animal bones, brownish-grey vessel with medium silt, slightly curved rim, split in the upper two-thirds, thick, wide wavy line decoration in the upper part; dated to the 8th century CE. Genetic sex: Male; Juvenis age.

#### 2 - Uniparental analyses

by Noémi Borbély, Veronika Csáky, Bea Szeifert

Great mtDNA diversity can be observed within the sampled pool of this study. Interestingly, contrary to the communities of the Great Hungarian Plain, the Eastern Eurasian lineages are less frequent in Transdanubia, although the typical haplogroups related to Hungarian conquerors (N1a1a1a1a, G2a1, *etc.*[10], [15], [22], [79]) and Avar elites (C4a1a4a, D4j5a, *etc.*[9], [10], [12]) are present (Fig. S5). There are several mtDNA haplotype identities within the sites of Himod, Zalavár and Sárbogárd which point to possible maternal kinship connections (for details, see Supplementary Information section 5). Between the two Himod horizons, no haplotype identities can be detected, though haplogroup identities can be. However, autosomal data indicate some level of continuity, as presented later and in the Main text. There are haplogroup level matches between the three horizons of the Zalavár cemetery, however, based on complete mitochondrial sequences, no haplotype similarities or identities were detected between these two groups, suggesting that there is no direct maternal relationship between the individuals of the different studied phases. According to the analyzed 56 Y-STR profiles, there are no identical Y chromosomes within the dataset compared to the mtDNAs.

### 3 - General genomic composition

by Bea Szeifert, Dániel Gerber

#### 3.1 PCA and Admixture

By organizing the studied groups from Transdanubian sites into century-wise batches for PCA analysis, we reveal a complex and eventful population history throughout the first millennium CE. The majority of the samples distributed on PC1 form a genetic cline in the region, later confirmed by further tests (Supplementary Information section 3.2) but Eastern Eurasian elements (negative values on PC2) can also be observed (Figures S6-S10). In the figures, Transdanubian samples marked with an *x* represent sampled, but chronologically irrelevant samples, on the corresponding figure. PCA backgrounds were compiled according to Jeong et al.[80] and Lazaridis et al.[81].

Population shifts and successions, discussed in the Main text and Supplementary Information sections 3.2 and 4, are summarized here based on the PCA (Figures S6-S10 and S12) and *ADMIXTURE* results (Fig. S11). The Székesfehérvár area and Himod face population shifts between the 8th and 10th centuries CE, likely connected to the Hungarian conquest (see Main text) but, very limited, population continuity can also be observed. On the contrary, Zalavár shows local succession throughout the centuries not only in genomic composition (PCA) but on an individual level (IBD and uniparental matches, see Main text and Supplementary Information sections 2 and 4) as well.

#### 3.2 *f*-statistics

We performed several $f_4$-statistics to distinguish individuals with and without East-Eurasian, Caucasian and/or Near Eastern ancestries in the form of $f_4$(Test, European, Eurasian, Mbuti.DG), where we selected a representative set of modern European populations and a representative set for all tested Eurasian regions through modern samples (*e.g.* East Asia, Siberia, for the complete set, see Table S3.1). We selected modern reference populations for this test particularly to avoid database bias caused by regional underrepresentation and variations in laboratory approaches used in different batches of ancient datasets[42]. Results are provided in Table S3.2. We also checked the results based on *ADMIXTURE* proportions, and found a correlation (Fig. S13), however, *ADMIXTURE* turned out to be inefficient to confidently exclude non-European ancestry, therefore we used only the $f_4$ results for the final classification.

To describe the genetic composition of the individuals with presumably only European ancestry, based on the previous $f_4$ tests, we conducted three separate $f_4$ in the form of $f_4$(reference_1, reference_2, Test, Mbuti.DG), where references were pairs of populations from Antonio et al.[82] and Gnecchi-Ruscone et al.[9] (see Table S4.1) covering distant regions of Late-Antique and Early Medieval Europe both geographically and genetically, for results, see Table S4.2. Then, we applied a clustering algorithm similar to Gerber et al.[35], which revealed three major groups (1-3), where Group 3 further subdivided into two subgroups (Groups 3.1 and 3.2, Fig. S14).

The distinction roughly follows the "European core" (referred to as CB-EUR here) established by Maróti et al.[10], although our approach simplifies these core groups of the Carpathian Basin. Groups 1 and 2 also have strikingly different uniparental compositions, as the former is mostly characterized by Y-chromosomal haplogroups E1b and J2, while the latter by haplogroups R1 and I (Table S1, Fig. S21). On the other hand, the uniparental composition and autosomal PCA intermediate position of Group 3 between Groups 1 and 2 (Table S5, Fig. 2) suggest that Group 3 was either 1) formed by the mixture of the former two, thus represents a natural genetic cline, or 2) as a "standalone" group, had major shared drift with both of them. We also performed two separate runs in the form of $f_4$(Group 1, Group 2, Test, Siberia) and $f_4$(Group 3.1, Group 3.2, Test, Siberia) on the remaining individuals with significant non-European ancestry, and two other in the same setting, but with the individuals of CB-EUR as tests, to test the aforementioned scenarios. For the $f_4$ results, see Table S4.3. The plotted results (Fig. S15) show a very strong attraction between Group 1 and 3.1, as well as between Group 2 and 3.2, respectively, suggesting that Group 3 is generally a cline between Groups 1 and 2, thus we hereafter refer to it as Cline 3. A conspicuous phenomenon, seen on Fig. S15 is that some samples are off from the highlighted "cline zone", which is likely due to not tested or hidden genetic relatedness. Notably, in the cladogram (Fig. S14), Group 2 is more closely attracted to Cline 3 compared to Group 1, which is likely due to the higher admixture rate of Group 2 into Cline 3, compared to Group 1.

Another striking feature can be seen in Fig. S16 of the East-Eurasian admixed individuals, where the pre- and post-CEE individuals form two distinct groups: the post-CEE individuals mostly overlap with contemporaneous CB-EUR Clines 2 and 3.2, whereas pre-CEE individuals form a rather distinct group around the position of Cline 3.1 (which is way less pronounced contemporarily) and in a non-cline dimension, foreshadowing a different West-Eurasian ancestry for this group, further discussed in the Main text. For individual group classification, see Table S5. As described in the Main text, a geographical pattern of these clines, even without the chronological distribution, can be observed when putting the classified individuals into a map (Fig. S17 and S18).

For the IBD analysis, we performed an $f_4$ in the form of $f_4$(Test, Asian, European, Mbuti.DG) (see Tables S4) to identify individuals without any West Eurasian ancestry. This test revealed a few individuals with non-European genetic ancestry.

### 4 - IBD analyses

by Kristóf Jakab, György Mező, Dániel Gerber

During the creation of Fig. 5 (“IBD patterns across Transdanubian sites across centuries”) in the Main text, we considered several archaeological and ecological features of the corresponding sites. Although Zalavár is only a few kilometers away from the sites of Vörs-Papkert, Hács and Szólád, it was treated as a separate region due to the marshland of Kis-Balaton, which even today acts as a strong ecological barrier. Accordingly, the Zalavár area is ecologically more similar to the Little Hungarian Plain (Himod, Árpás and Szakony sites), however was treated separately due to their greater distance. The site of Vörs-Papkert was also considered separately for its ecologically transitional position between Zalavár and the other Southeastern Transdanubian sites (Szólád, Hács, Fonyód, Balatonszemes, Bodajk, Székesfehérvár, Pákozd, Csákberény, Előszállás).

### 5 - Genetic relatedness analyses

by Veronika Csáky, Bea Szeifert

We mapped several first- and second-degree biological relationships within our collected samples using *READ*[40] and *KIN*[41]. In Himod I, KOZ03 (Grave 127, 7-9 year old girl) and KOZ12 (Grave 128, 30-39 year old female) are first-degree relatives, likely mother and child, as they are in adjacent graves and their mitochondrial DNA matches (U3b2a1). Another first-degree relatedness in Himod I is between two sisters (KOZ05, Grave 153, 10-14 years old and KOZ10, Grave 151, 0.5-1.5 years old), also with matching mitochondrial DNA (H13b1). A third sample (KOZ01, Grave 104, 9-12 year old girl) shares the same mtDNA (H13b1) with the above mentioned individuals, but without full genome analysis, the exact relationship is indeterminate. They share a common maternal ancestor, as indicated by the proximity of their graves in the cemetery. We found no kinship connections among the individuals of the Himod phase II and between the two different phases.

In Sárbogárd, two boys (AHP143, Grave 15, 6 years old and AHP144, Grave 23, 3 years old) are brothers, with the same mtDNA (J1b1a1) and Y-haplogroup (R1b1a1b1a1a2c1). An older female (AHP142, Grave 10, 62-75 years old) also shares their mtDNA (J1b1a1), suggesting a maternal relationship, but without autosomal data, the exact link remains unknown. All three graves are part of the late group of the cemetery, although the older female's grave is slightly further from the boys'. Additional relationships in Sárbogárd include a second-degree relationship between AHP139 (Grave 72, 48-52 year old male) and AHP154 (Grave 78, 17-18 year old male), and a second or third-degree relationship between AHP152 (Grave 74, 32-36 year old male) and AHP153 (Grave 75, 40-48 year old male). These individuals have different mitochondrial haplogroups, excluding kinship through direct maternal lineages. The second-degree relatives belong to different subgroups of the I1a2a2 Y-chromosomal haplogroup, thus they are likely maternal uncle-nephew or maternal grandfather-grandson relatives. The third-degree relatives share the same Y-haplogroup (R1b-L20), thus paternal cousins can be assumed but other scenarios are also possible. Both of these samples are part of the late group in the cemetery, located next to each other.

At the Bodajk – Homoki-dűlő site, we identified two first-degree relationships: AHP204 (Grave 4, 50-54 year old male) and AHP206 (Grave 19, 30-39 year old male) are brothers, sharing both maternal and paternal lines (T2b5a, E1b1b1a1b1a), and another between AHP207 (Grave 23, 40-44 year old male) and AHP208 (Grave 40, age 40-x female) as parent-child, specifically mother-son, with matching mitochondrial lineage (D4j8).

At Bodajk – Proletárföldek, AHP190 (Grave 2, male) and AHPS195 (Grave 11, 3-4 year old boy) are first-degree relatives, father and son, as they have different mitochondrial lines but the same Y haplogroup (R1a1a1b1a1a1c1l~). A second-degree maternal relatedness (H6c, differing Y line) was detected between AHPS194 (Grave 8, male) and AHPS202 (Grave 27, 30-34 year old male). For detailed results, see Tables S6 and S7.

Identical mtDNA haplotypes between samples may indicate maternal relatedness or population continuity across horizons, and could provide some additional information when autosomal data is missing. In Table S9 we list identical maternal haplotypes that suggest several connections within the Himod, Sárbogárd and Visegrád sites but no such connections within Zalavár were observed.

Further connections detected between the Transdanubian samples and the analyzed published genomes are discussed in the Main text. All connections are presented in Table S7. Distant connections mapped via IBD are visualized in Fig. S26.

### Supplementary Figures

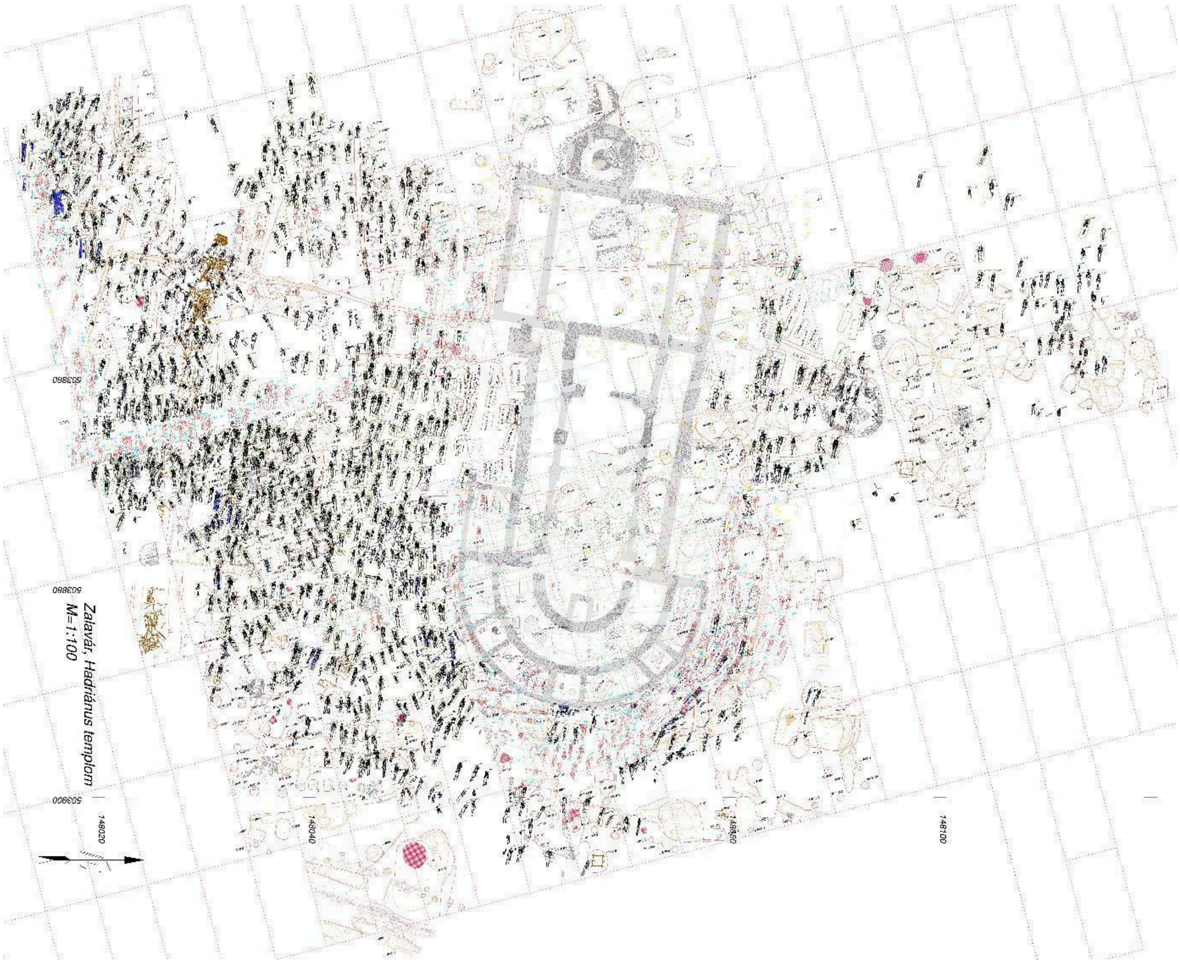


*Figure S1: Burials around Hadrian's temple at the Zalavár-Vársziget site (9th-12th centuries CE).*

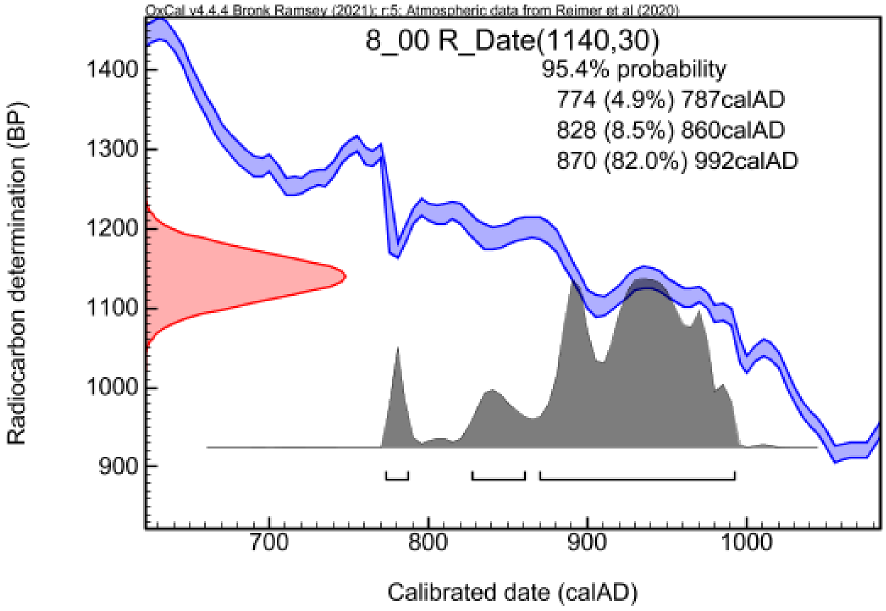

***Figure S2: Calibrated radiocarbon date and curve of the individual from grave 8/00 at Zalavár.***

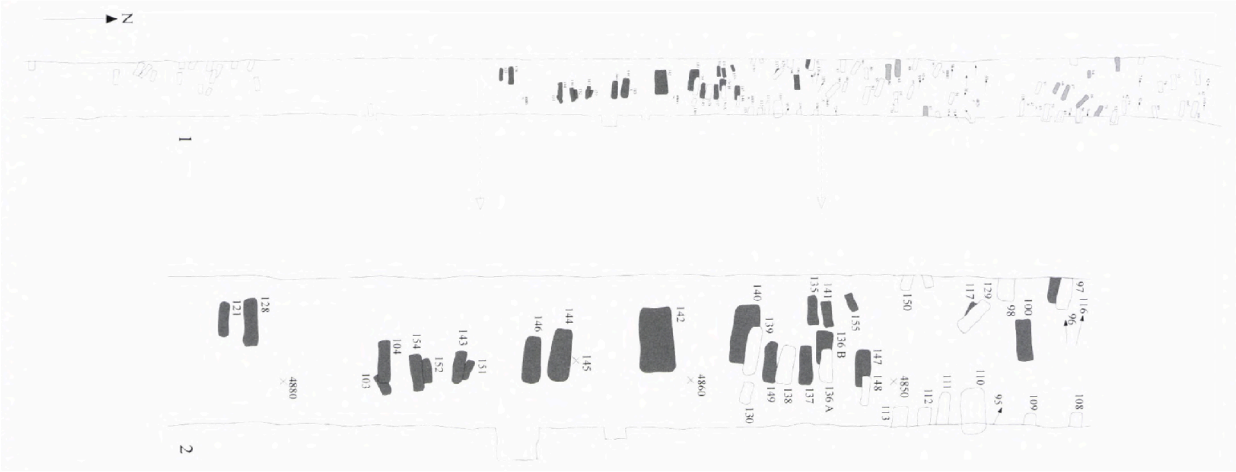

***Figure S3: Himod - Káposztásföldek: showing the 9th century CE graves in dark grey, and the 10th-11th century CE horizon in light grey[51].***

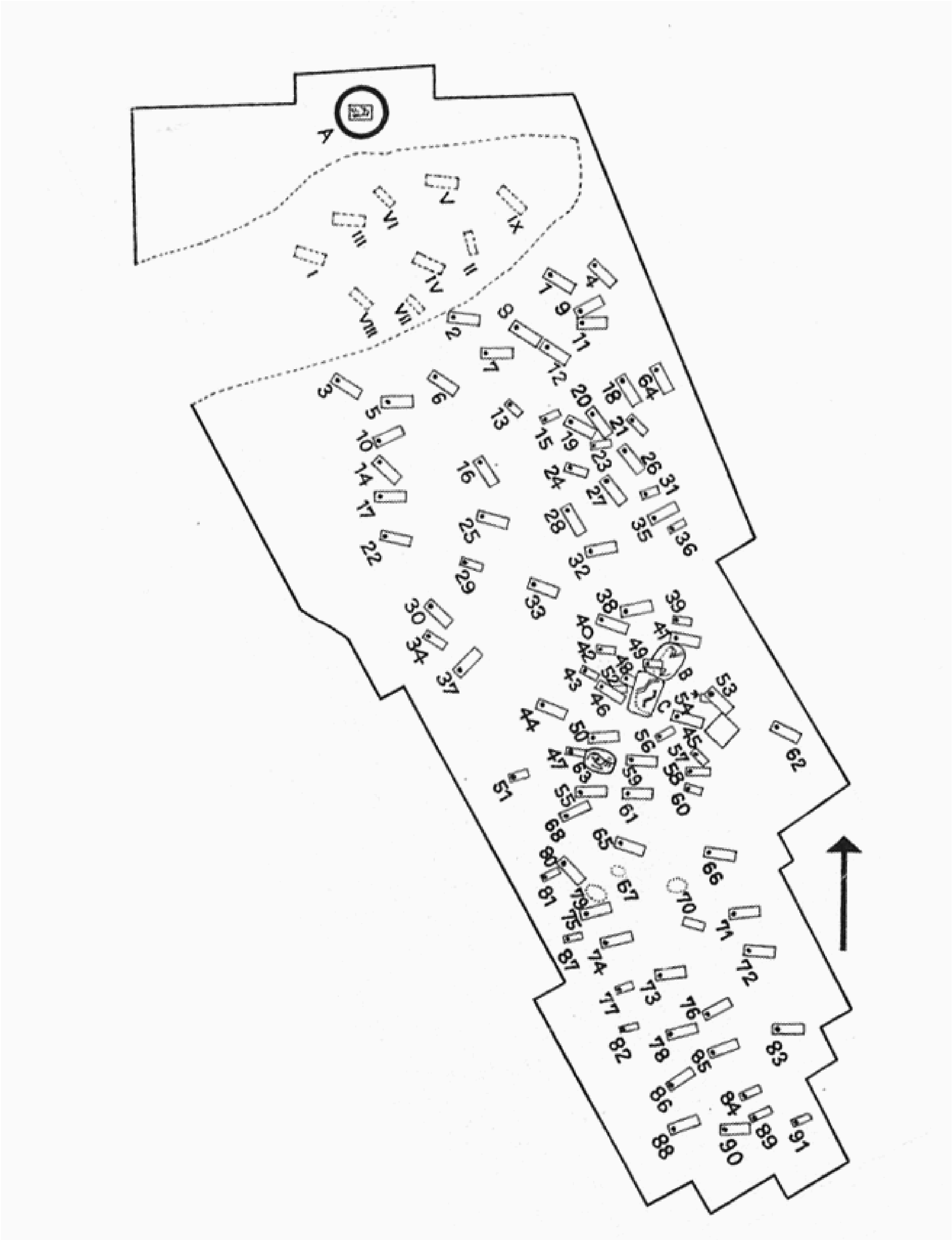

***Figure S4 Location of the graves at the Sárbogárd-Tringer tanya site following Éry 1967.***

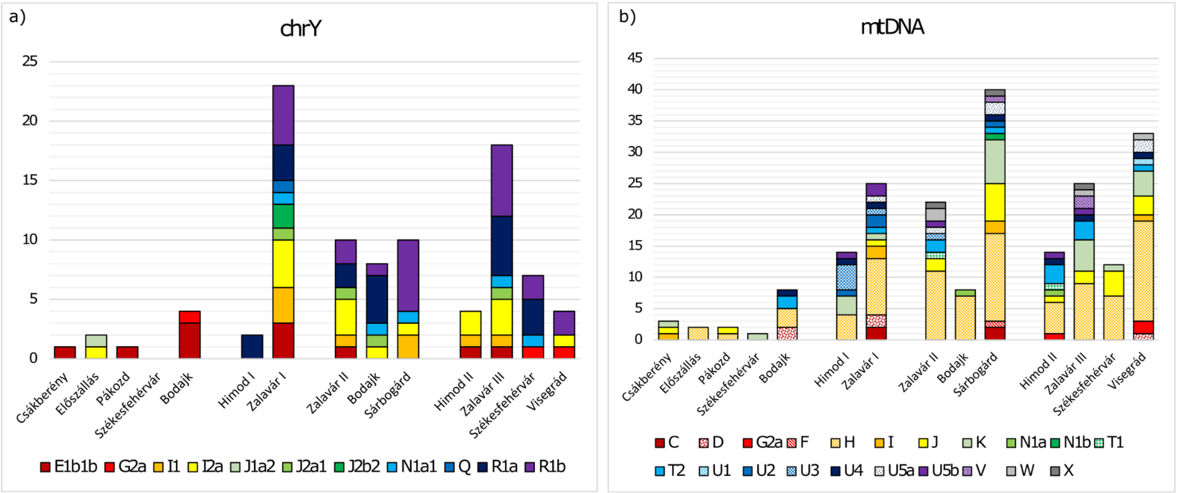


***Figure S5. Uniparental composition of the studied sites***

*a) Y-chromosomal composition of the studied sites. b) MtDNA (mitochondrial DNA) composition of the studied sites. X axis corresponds to chronology and Y axis to sample number in both figures.*

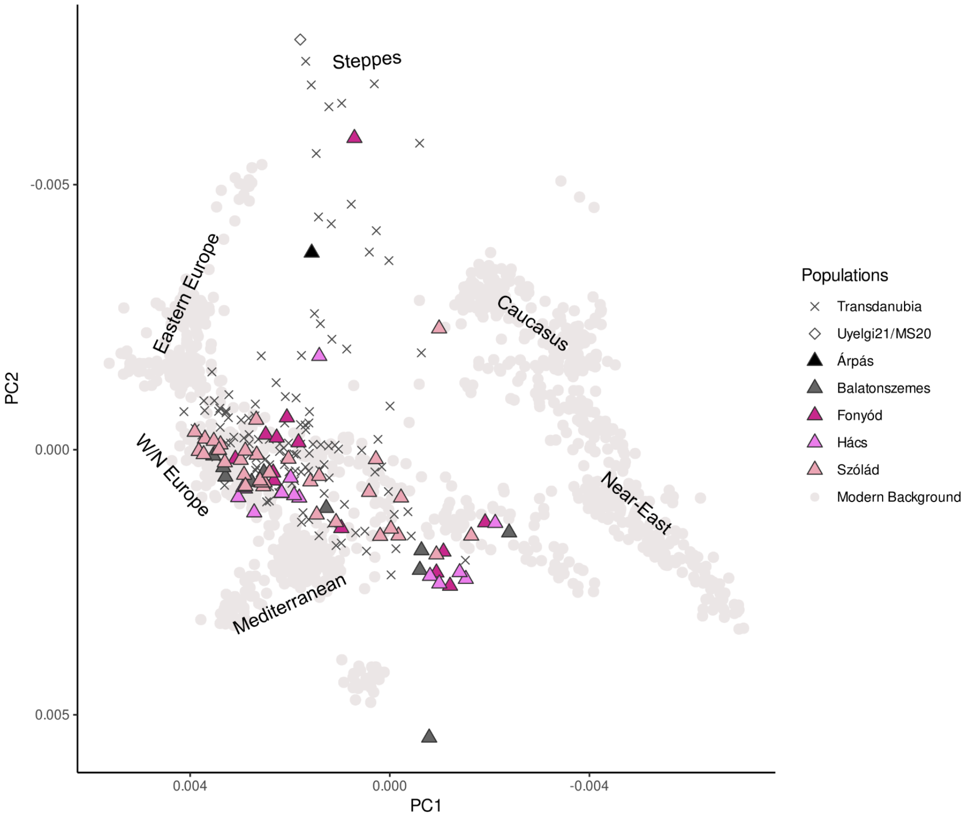


***Figure S6. Principal component analysis (PCA) of Transdanubian individuals in the 5th-6th centuries CE***

*Individuals before the 6th century CE mostly from Amorim et al.[8] and Vyas et al.[11].*

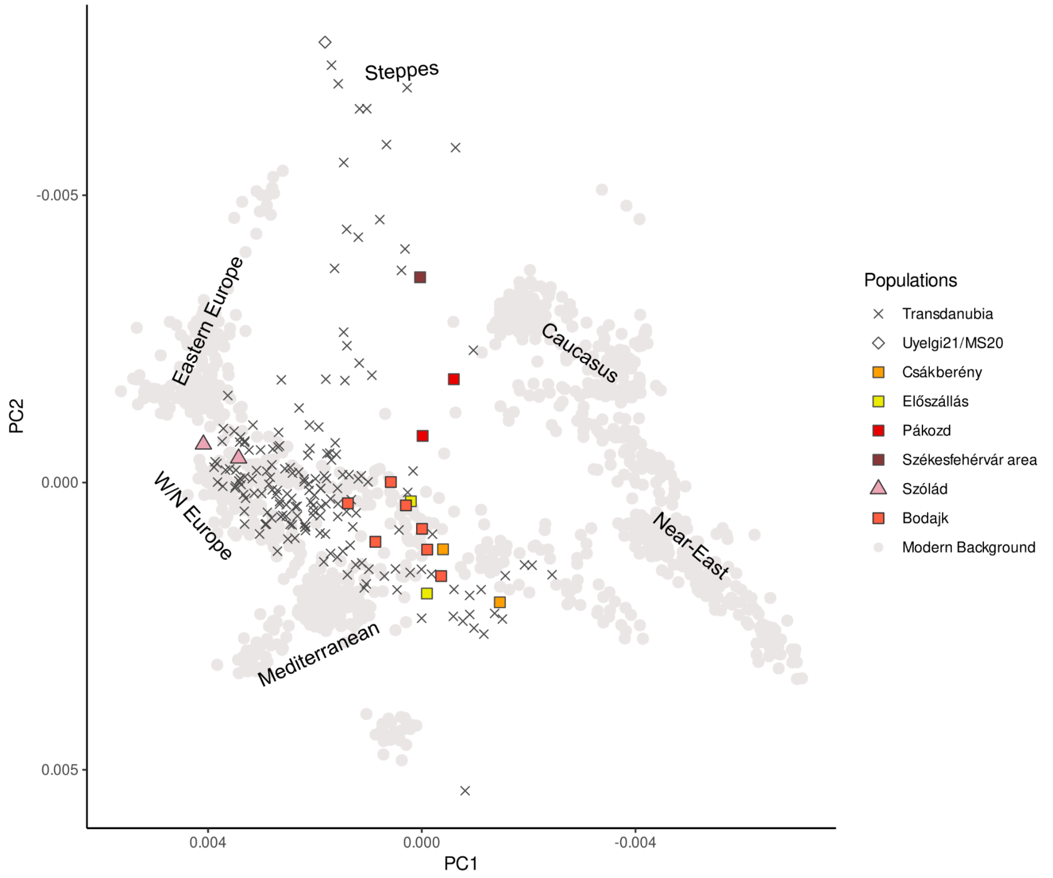


***Figure S7. Principal component analysis (PCA) of Transdanubian genomes from the 7th-8th centuries CE***

*A major change in the genomic makeup after the 6th century CE can be observed among the collected samples from Székesfehérvár area and proximal sites (Csákberény, Előszállás, Pákozd, Bodajk). There are almost no samples with positive first principal component (PC1) values, suggesting the presence of a rather southern European characteristic base population, except individuals AV1 and AV2 from Szólád. Additionally, several samples with eastern ancestry appear that can be connected to the pre-CEE cluster (see Main text). AV1 and AV2, while overlapping on the PCA with samples from the former centuries, represent a shift in ancestry that aligns with discontinuity in the IBD (identity-by-descent) connection system (Supplementary Information section 4, Main text).*

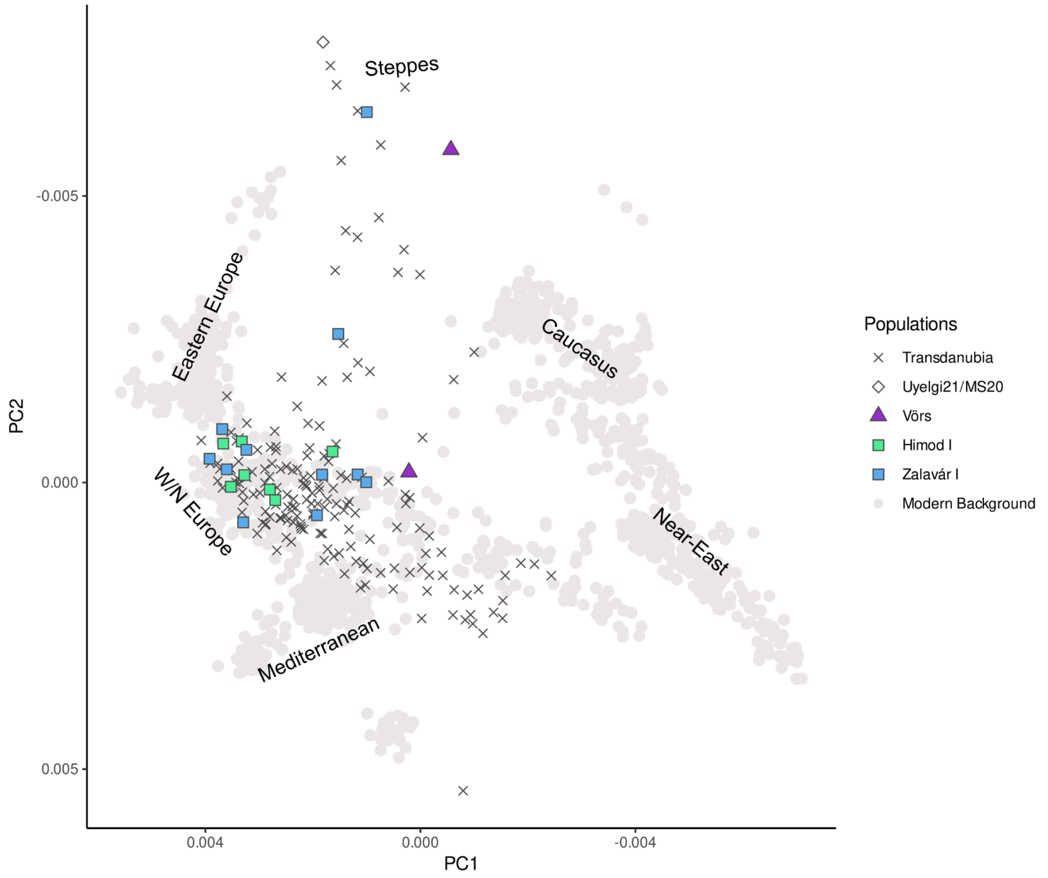


***Figure S8. Principal component analysis (PCA) of Transdanubian genomes from the 9th century CE***

*The 9th century CE phase mostly consists of Zalavár I and Himod I samples, thus the apparent shift in genomic makeup could be at least partially aligned with differences between people living in different ecoregions (detailed in the Main text). The majority of the 9th century CE individuals overlap with individuals AV1 and AV2 from the 7th century CE on the PCA due to common ancestry, also highlighting the importance of a relatively hidden substrate between the 7th-8th centuries CE (see Main text, Supplementary Information section 4). Individual AHS21 is highly dissimilar to the majority of the samples plotted on the PCA and likely represents one of the earliest forerunners of Hungarian conquerors, as later confirmed by ADMIXTURE (Fig. S11) and IBD (identity-by-descent) analysis (Main text).*

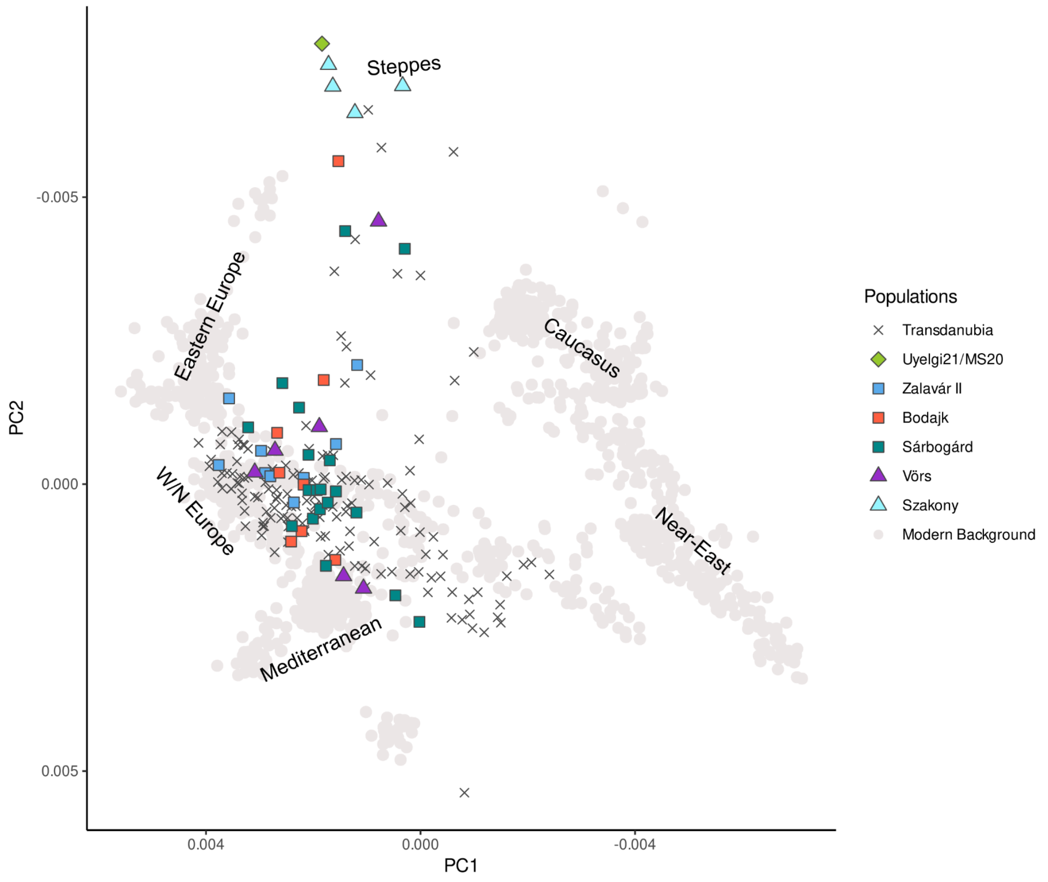


***Figure S9. Principal component analysis (PCA) of Transdanubian genomes from the 10th century CE***

*The PCA of 10th century CE Transdanubia shows no major shift in ancestries, although a greater sample dispersion and the lack of negative first principal component (PC1) value samples indicate the dilution or disappearance of southern European genetic elements in proximal sites of the Székesfehérvár area (Sárbogárd, Bodajk, Main text, Supplementary Information section 3.2 and 4) compared to Fig. S7. The extensive appearance of Hungarian conquerors is also apparent in this horizon, such as the Szakony individuals in close proximity to individual Uyelgi21/MS20 (Ural region).*

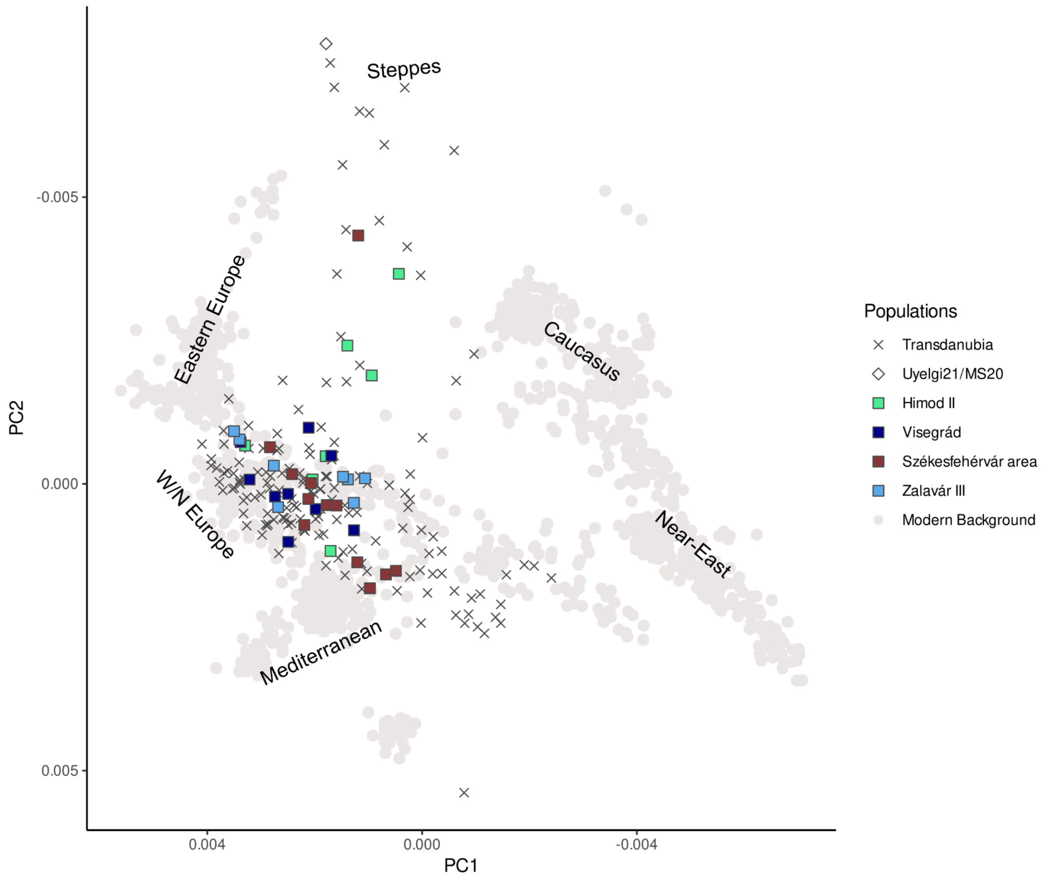


***Figure S10. Principal component analysis (PCA) of Transdanubian genomes from the 11th century CE***

*After the 10th century CE, a homogenisation process can be observed in the Carpathian Basin that was confirmed by other analyses (Main text, Supplementary Information section 3.2 and 4).*

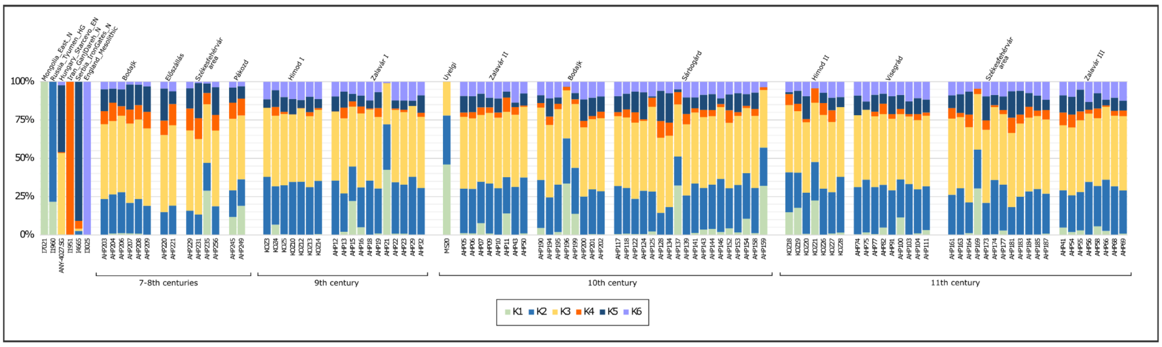


***Figure S11. K=6 unsupervised admixture of Transdanubian individuals and the reference genome Uyelgi21/MS20 from the 8th-10th century CE Trans-Ural region.***

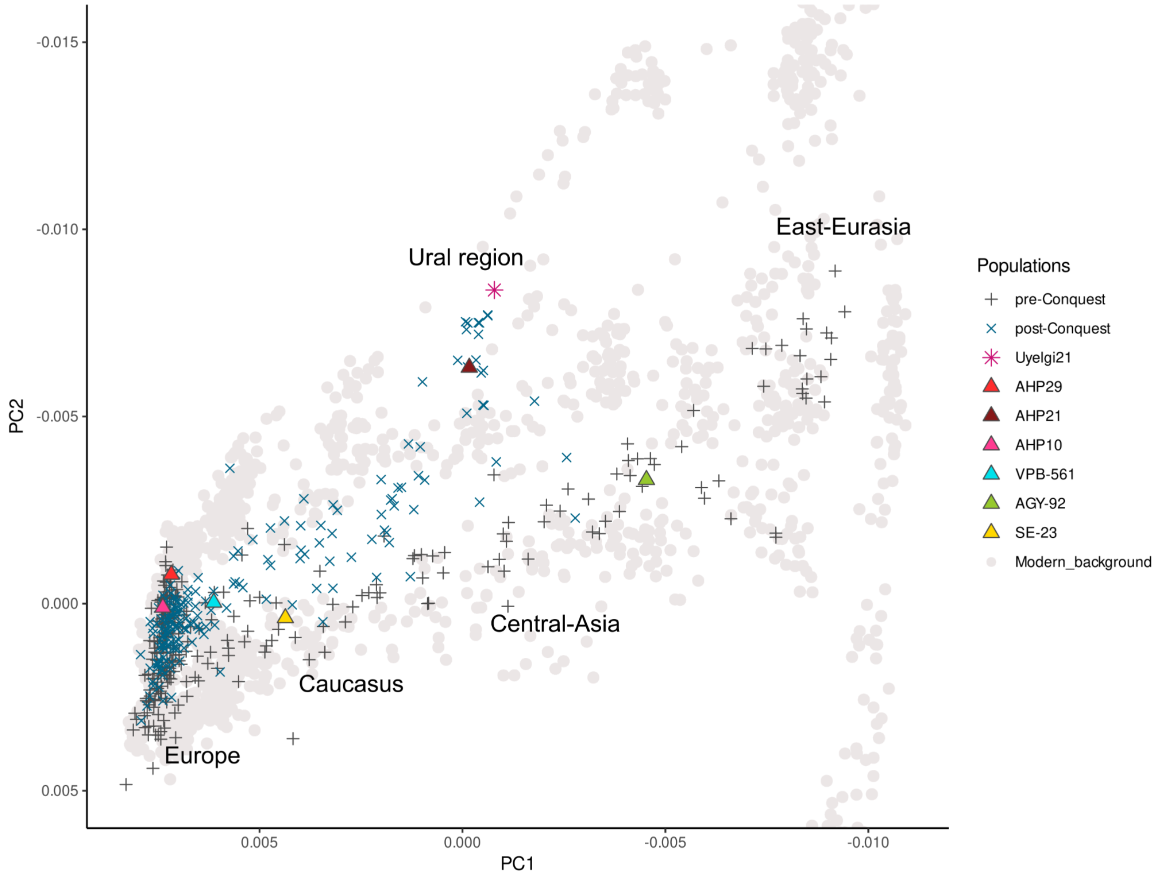


***Figure S12. Principal component analysis (PCA) with Eurasian background***

*All studied individuals are projected onto the modern Eurasian genetic diversity, where we marked pre- and post-Conquest period individuals. The separation of the two groups reveals two PCA clines that are easily paralleled with the two IBD (identity-by-descent) clusters (see Main text). We highlighted those individuals whose chronological and IBD-cluster groupings differ: AHP21 and AHP29 come from the pre-Conquest period but belong to the Conqueror cluster. AHP10, VPB-561, AGY-92, SE-23 represent the 10th century, but were included in the Avar cluster during the analyses (Table S5).*

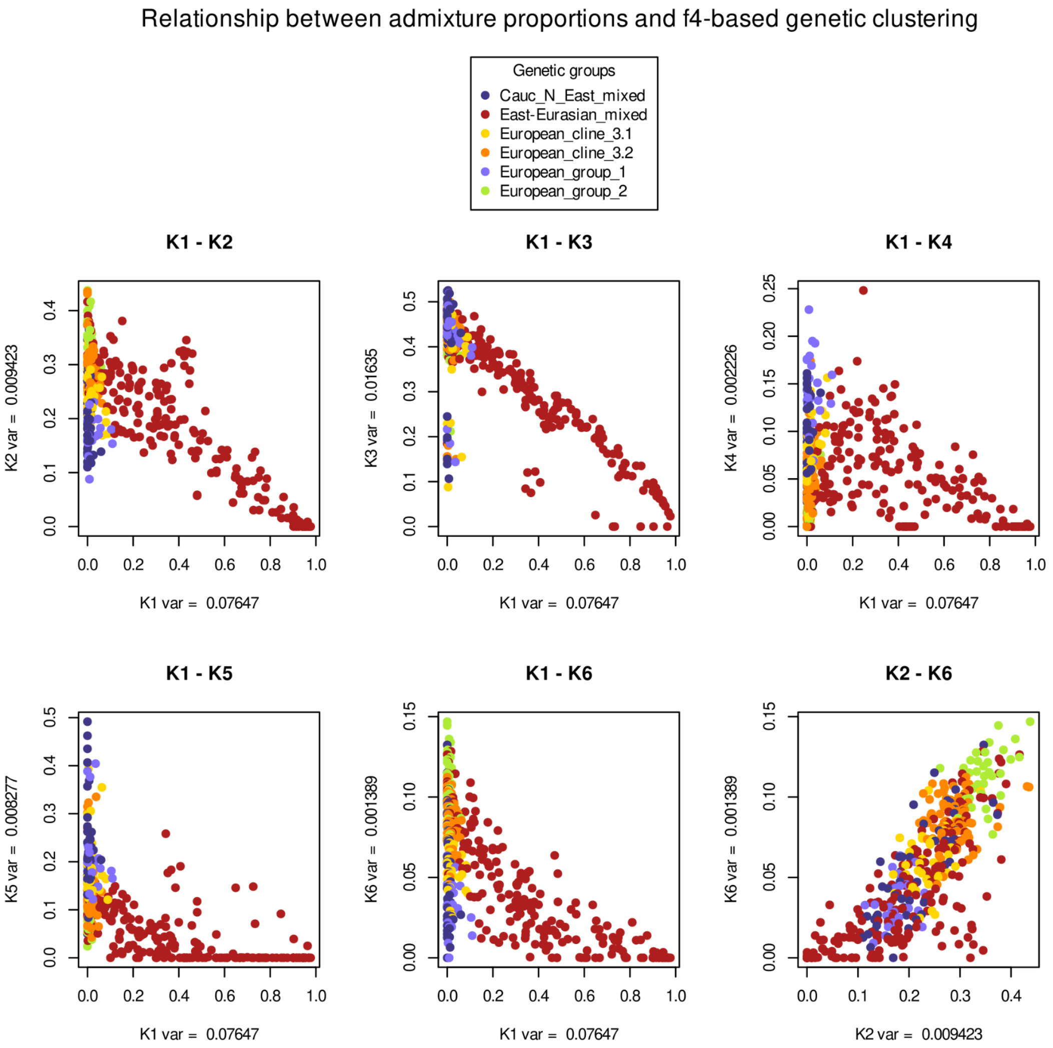


***Figure S13. Comparison of $f_4$- and ADMIXTURE-based classification of studied individuals.***

*As it is demonstrated on this figure, ADMIXTURE and f-statistics results correlate with each other, and highlight the validity of our novel $f_4$ statistics-based clustering and filtering approach.*

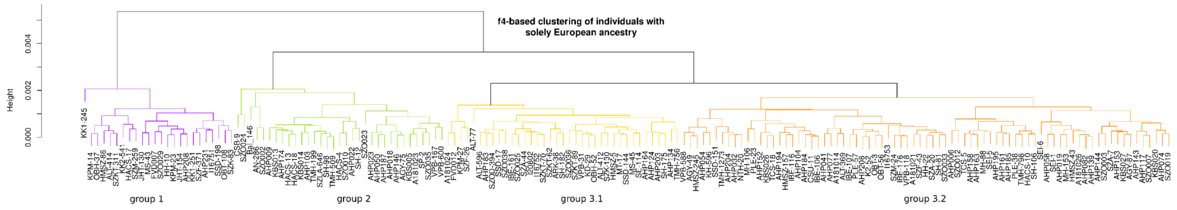


***Figure S14. $f_4$-based clustering (see Supplementary Information section 3.2) of individuals with European ancestry.***

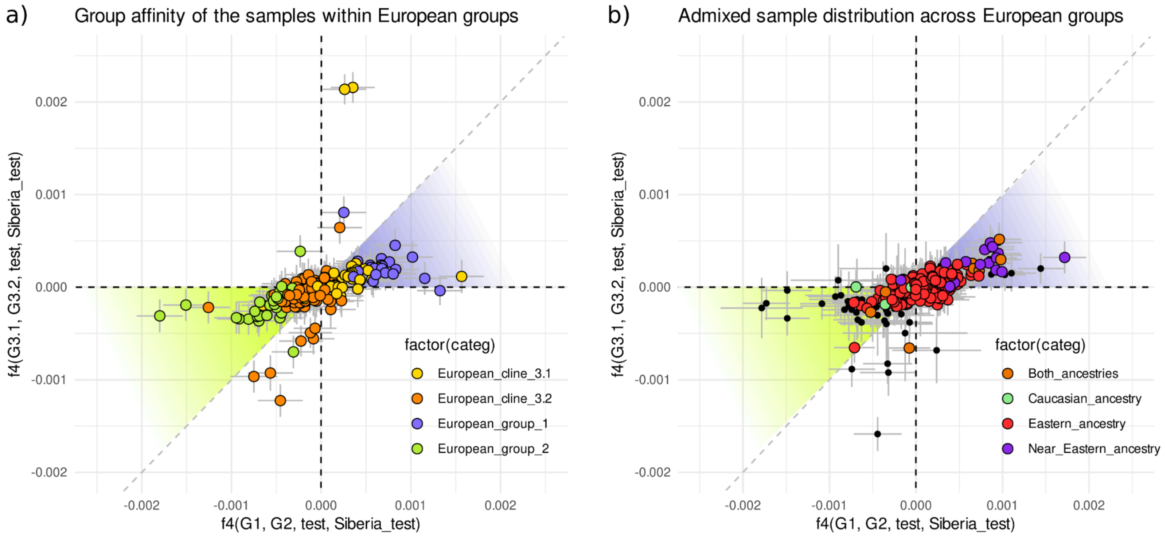


***Figure S15. Distribution and affinities of European groups/clines***

*Classification of a) CB-EUR and b) admixed and low coverage individuals into European genetic Groups 1-2 and Cline 3. Caucasian (green), Near Eastern (purple), a combination of these (orange), and East-Eurasian (red) ancestries are highlighted. Near Eastern ancestry is restricted to Group 1 and Cline 3.1 affinities, suggesting strong geographic clinality of the European population in the Carpathian Basin. The highlighted areas represent the threshold between Clines 3.1-1 and 3.2-2 affinities if Cline 3 is the two-way mixture of Groups 1 and 2. For the vast majority of samples this fits well, and for some samples, the position outside the highlighted area is mostly due to genetic relatedness between plotted samples and references used.*

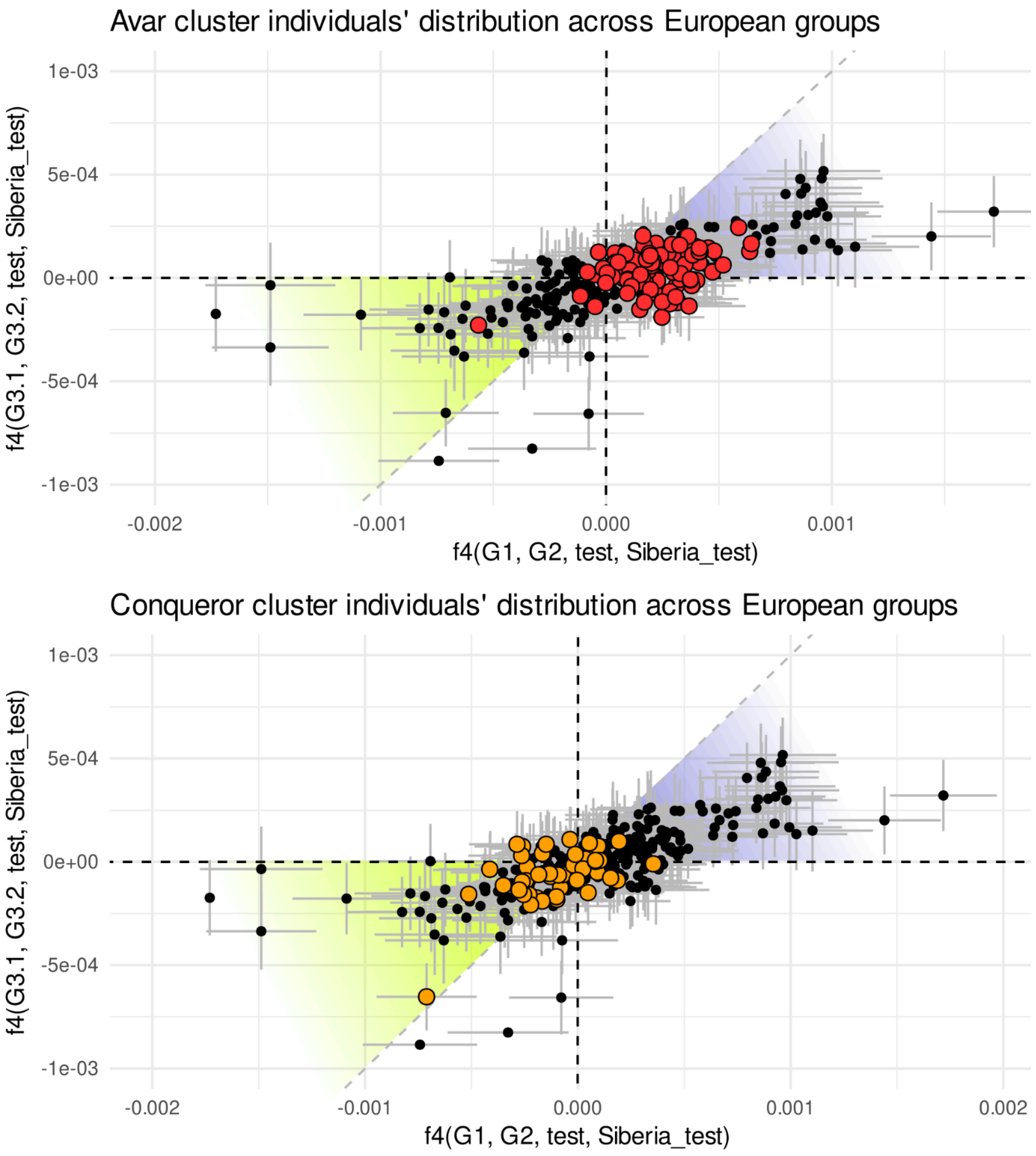


***Figure S16. Affinities of East-Eurasian admixed individuals to CB-EUR groups/clines.*** *Black dots represent all admixed/low coverage individuals, red dots the pre-CEE (Avar) cluster and orange the post-CEE (Conqueror) cluster individuals. There is only minimal overlap between Avar and Conqueror cluster individuals signaling an overall different West-Eurasian genetic background for the two groups.*

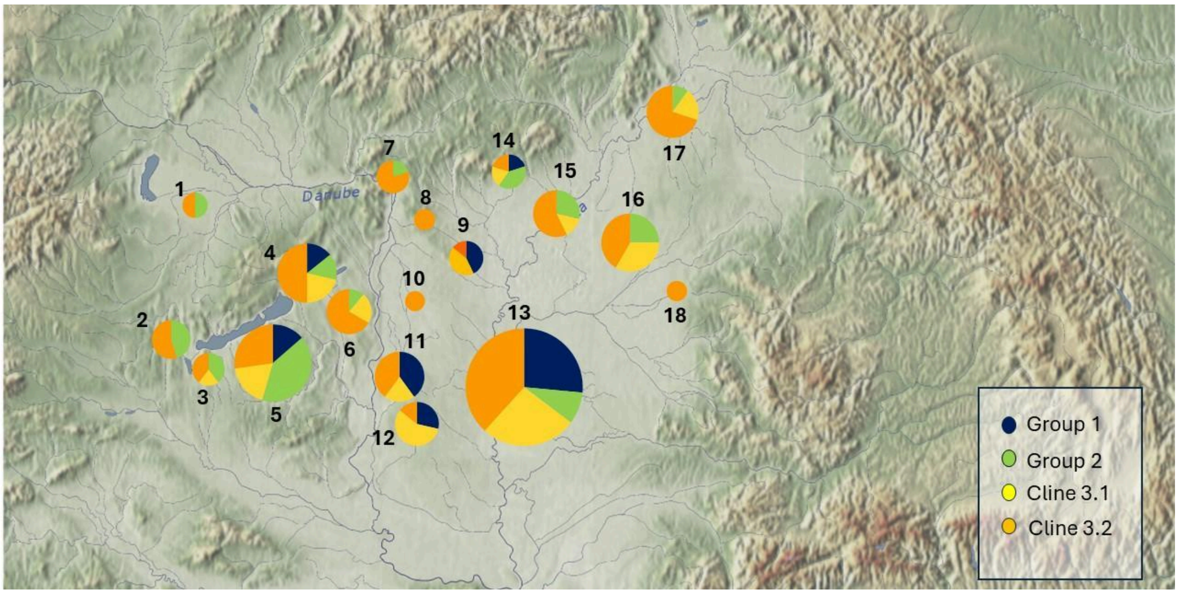


***Figure S17. Geographical distribution of CB-EUR individuals, coloured by their corresponding genetic cline.***

*1. Himod (n=6); 2. Zalavár (n=11); 3. Vörs-Papkert (n=5); 4. Bodajk, Csákbereny, Székesfehérvár (n=14); 5. Balatonszemes, Fonyód, Hács, Szólád (n=22); 6. Sárbogárd (n=9); 7. Visegrád (n=5); 8. Nagytarcsa (n=1); 9. Alattyán, Jánoshida (n=7); 10. Kecskemét (n=1); 11. Homokmégy, Kiskőrös (n=10); 12. Madaras, Mélykut, Sükösd (n=7); 13. Algyő, Apátfalva, Csongrád, Kiskundorozsma, Orosháza, Sándorfalva, Szegvár, Székkutas, Szeged (n=34); 14. Szilvásvárad, Visonta (n=5); 15. Tiszafüred, Tiszanána (n=7); 16. Árkus, Kaba, Püspökladány, Sárrétudvari (n=12); 17. Hajdúnánás, Ibrány-Esbó-Halom, Karos II (n=10); 18. Magyarhomoróg (n=3).*

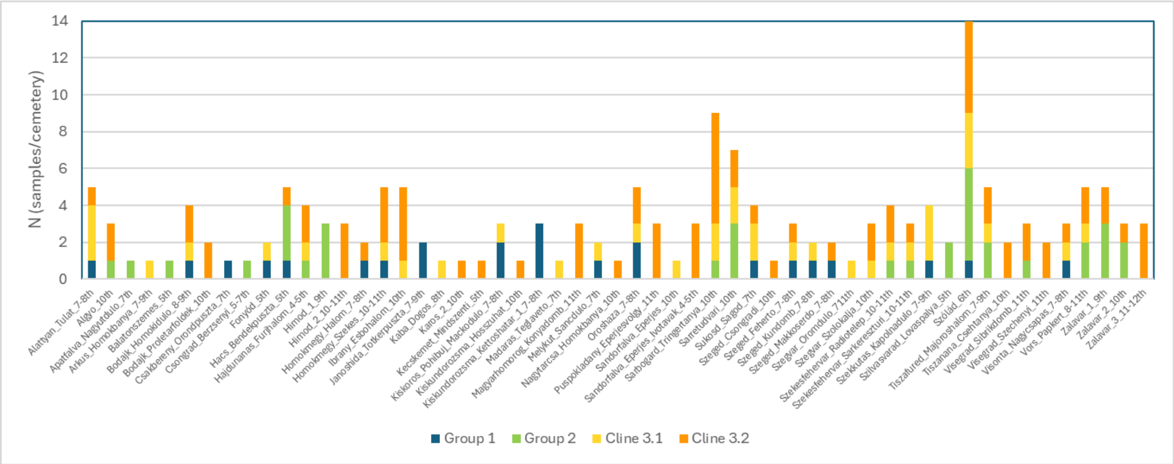


***Figure S18. Site-wise distribution of CB-EUR individuals, coloured by their corresponding genetic cline.*** *For details of the site-wise distribution, see Table S5.*

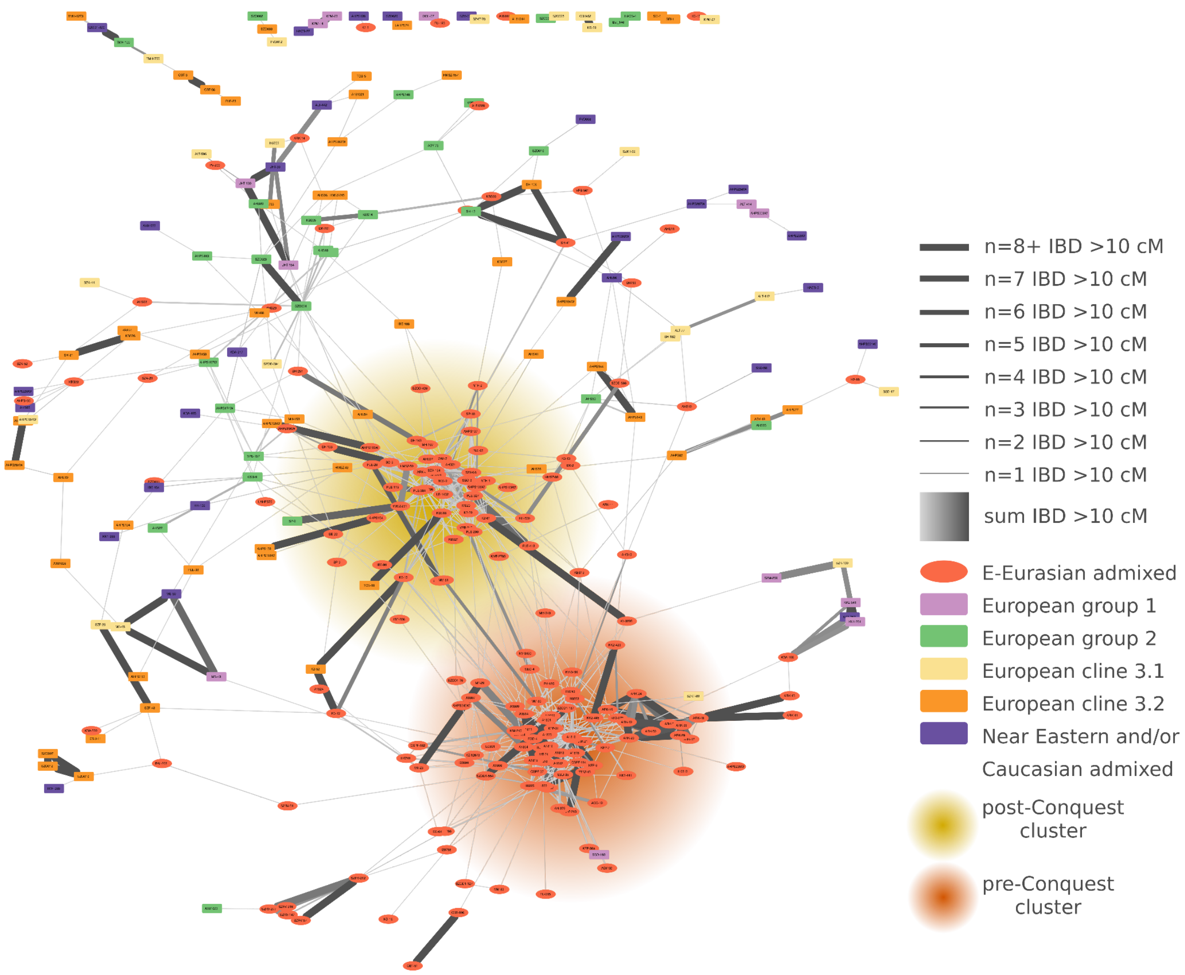


***Figure S19. IBD (identity-by-descent) network of >10 cM shared segments.***

*The edge-weighted spring embedded layout, driven by the calculated edge-betweenness (a measure to how often an edge lies on the shortest paths between pairs of nodes in a network). Individuals with no IBD connections are not shown. The lower dense group mostly consists of pre-9th century CE individuals with eastern ancestry, while the upper dense group mostly consists of post-9th century CE individuals with eastern ancestry. This is basically equal to eastern Avars and Hungarian conqueror genetic distribution.*

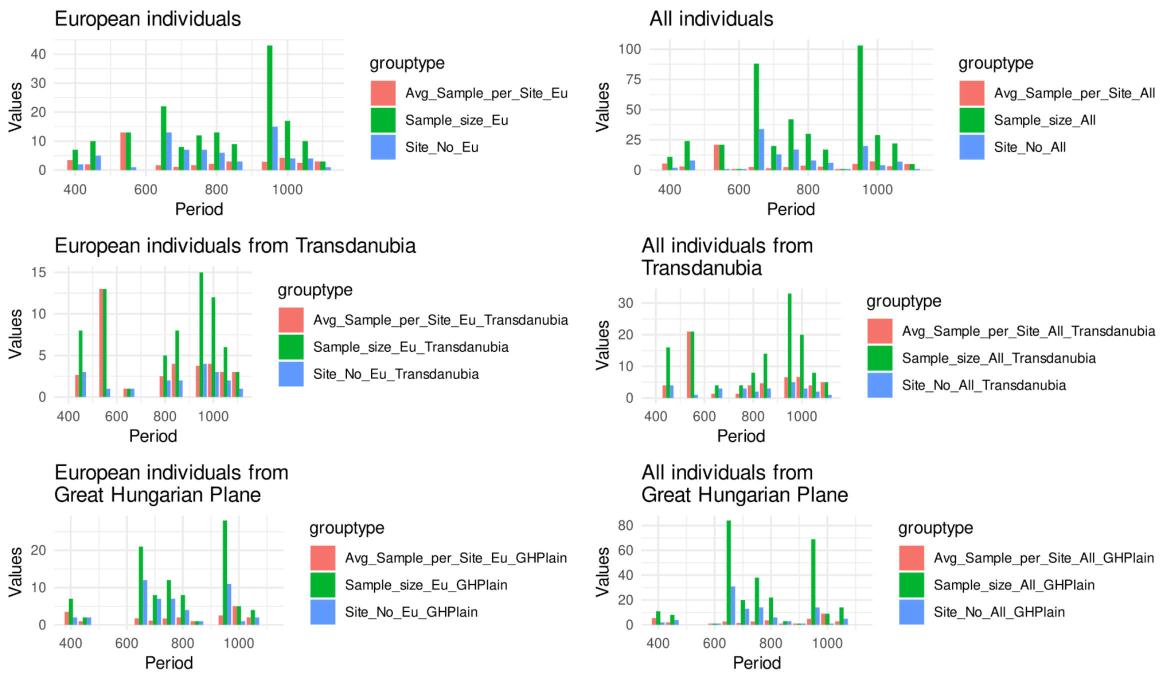


***Figure S20. Sample distribution across centuries and sites.***

*Red columns show the average sample per site values, green the sample size, and blue the site numbers of the correspondent centuries.*

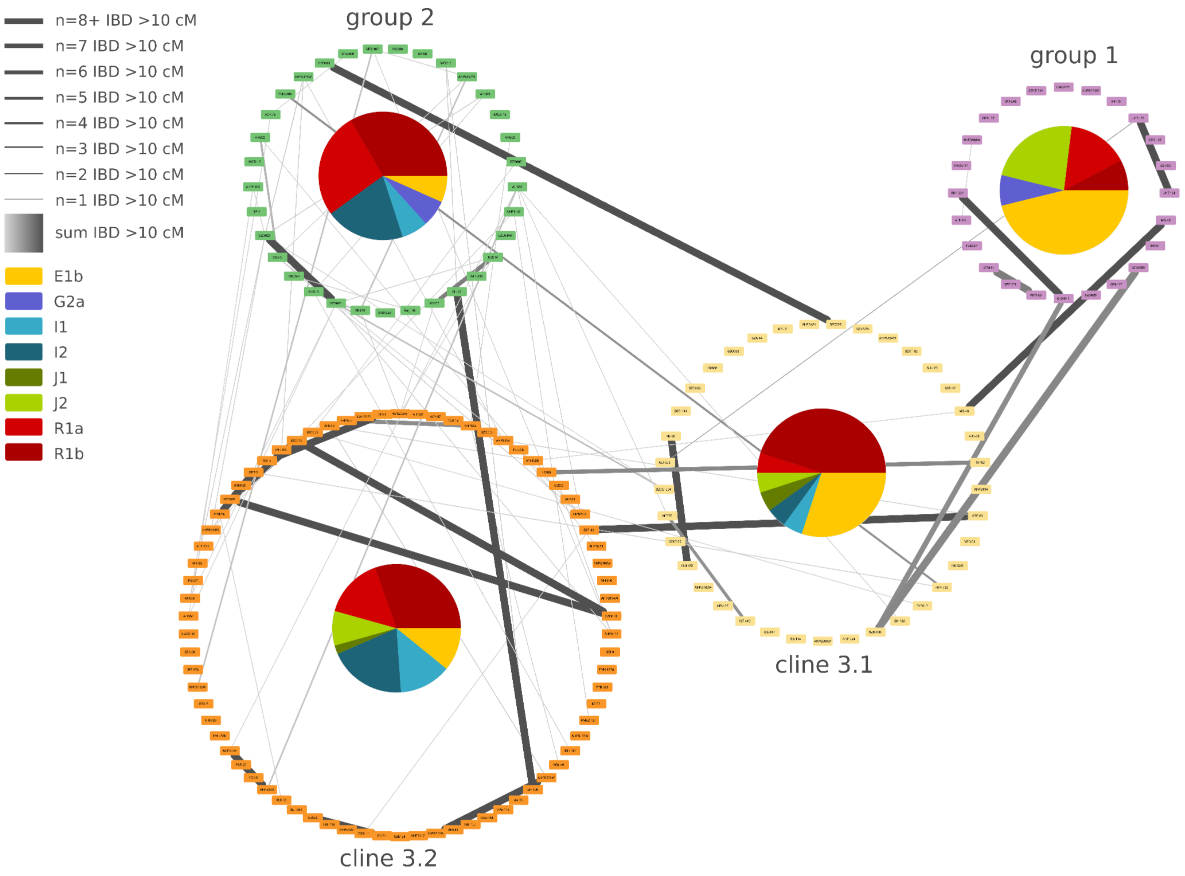

***Figure S21. IBD (identity-by-descent) chromosome segment sharing connections of European genetic Groups 1-2 and Cline 3 with Y-chromosomal haplogroup proportions.*** *The connections of the groups is in line with the $f_4$-predicted relationship system between the European core genetic groups (>10 cM (centimorgan) shared segment). Whereas 3.1 and 3.2 groups have intermediate positions in the connection system, Groups 1 and 2 have no IBD connections with each other at all.*

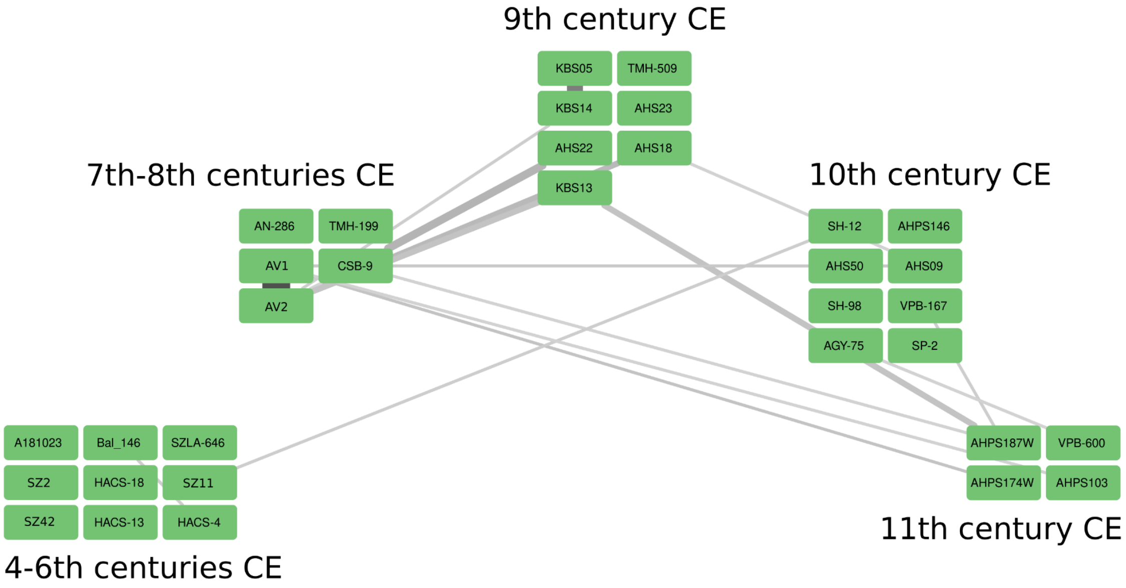


***Figure S22. IBD (identity-by-descent) chromosome segment sharing connections of European genetic Group 2.*** *Minimum 10cM segment sharings were considered. Discontinuity between the 6th-7th centuries CE is evident among individuals of the CB-EUR Group 2. This is also reflected by a shift in Y chromosomal haplogroups, as 4th-6th centuries CE individuals are mostly defined by R1b/I2a2, and the laters have R1a/I2a1 lineages within Group 2.*

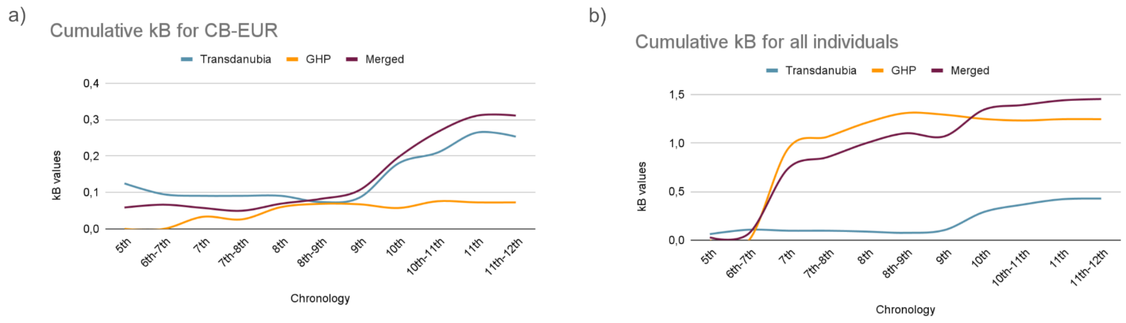


***Figure S23. Intergroup tendencies in IBD (identity-by-descent) segment sharing patterns using the 2n normalization method.*** *a) Cumulative connections between sites ($k_B$)*

*regarding the CB-EUR cluster in a chronological order. The formula used here balances sample size differences between sites and chronological batches, thus it highlights relative trends. Contrary to Fig. 4a, the trend for the Transdanubian communities show much more pronounced inter-site connections at the turn of the 9th-10th century CE, but there are no further major differences between further trends for those calculated with the ni*nx normalization (Fig. 4a and b, see Methods for the formulas).*

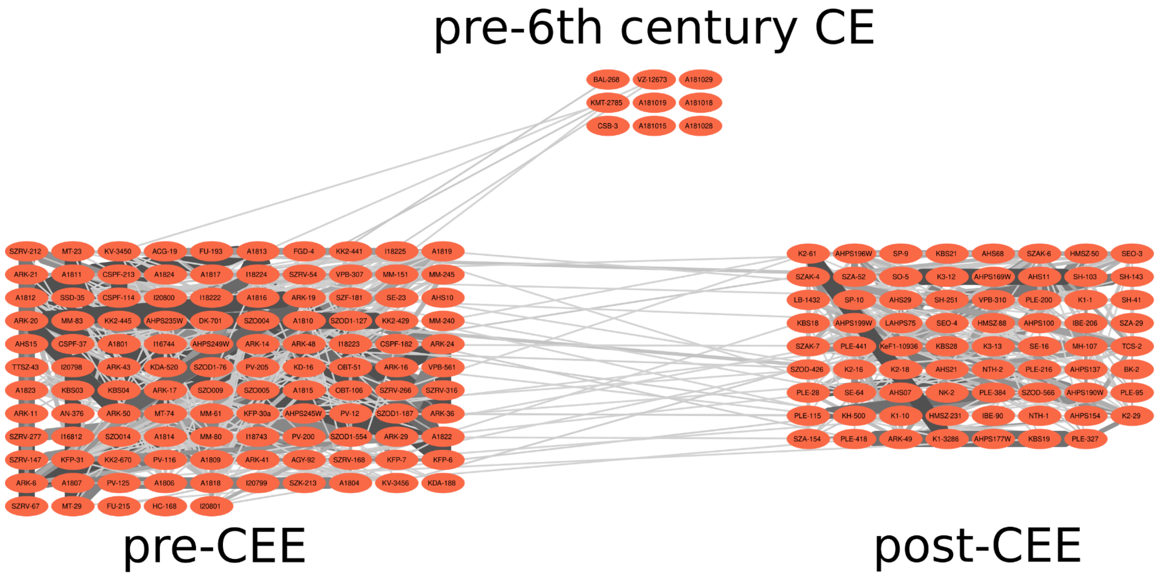


***Figure S24. IBD (identity-by-descent) chromosome segment sharing connections of individuals East-Eurasian ancestry***

*It is to observe the loss of pre-6th century East-Eurasian ancestry in the post-CEE cluster. Connections of pre-6th century individuals to the pre-CEE cluster could either indicate local succession and/or shared origins.*

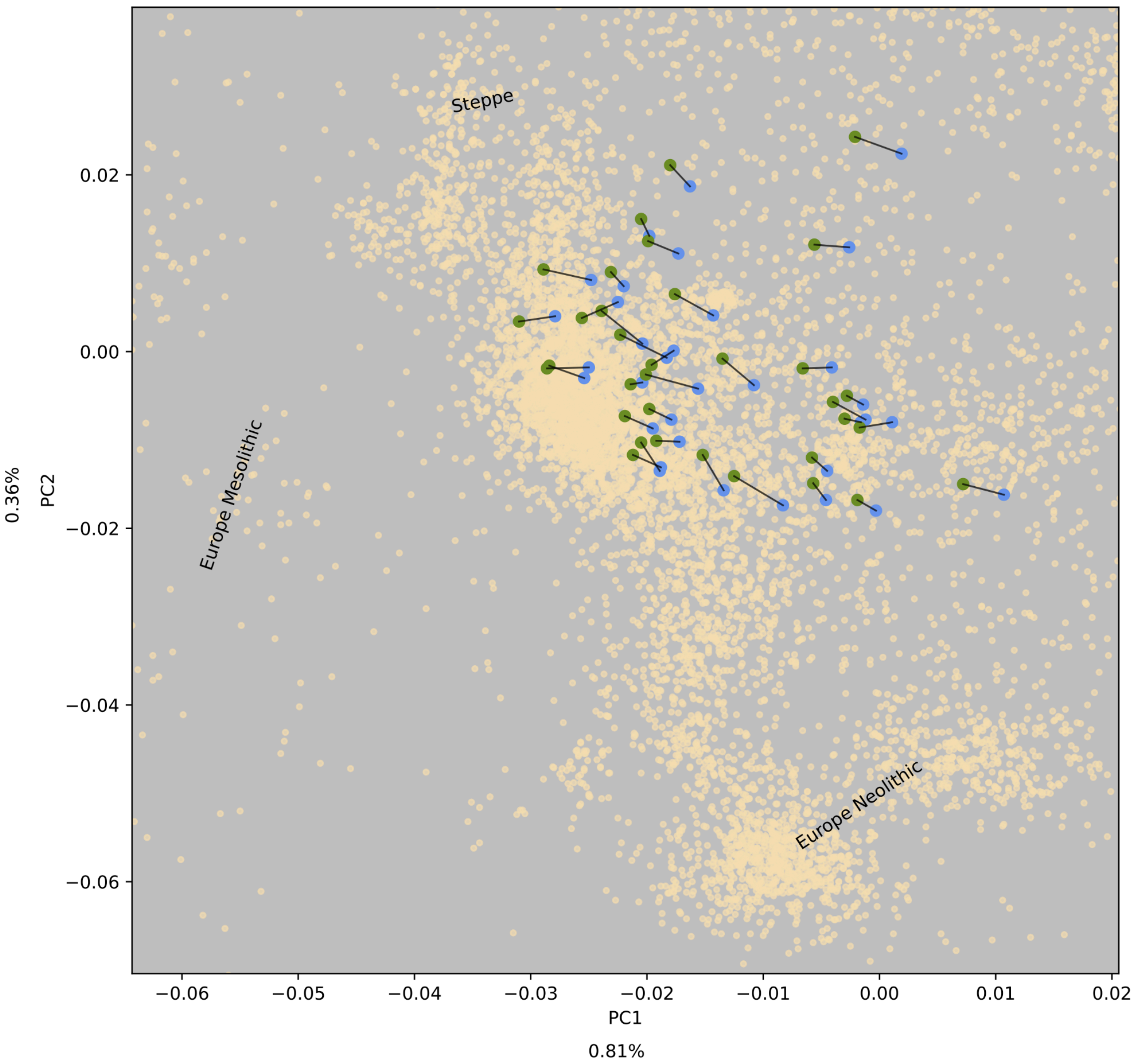


***Figure S25. Imputation accuracy***

*This plot demonstrates imputation accuracy of ancient genomes of this study by PCA (principal component analysis) calculation (with smartpca) and visualization. The original pseudo haploid genomes (blue dots) and their phased and imputed pairs (green dots) are plotted, connecting the identical samples with a black line. Unsurprisingly, most likely due to the asymmetric reference panel (1000 Genomes Project) used for imputation, most samples show a shift towards one direction, pointing out the biased nature of imputation, which could alter ancIBD results to a minor extent.*

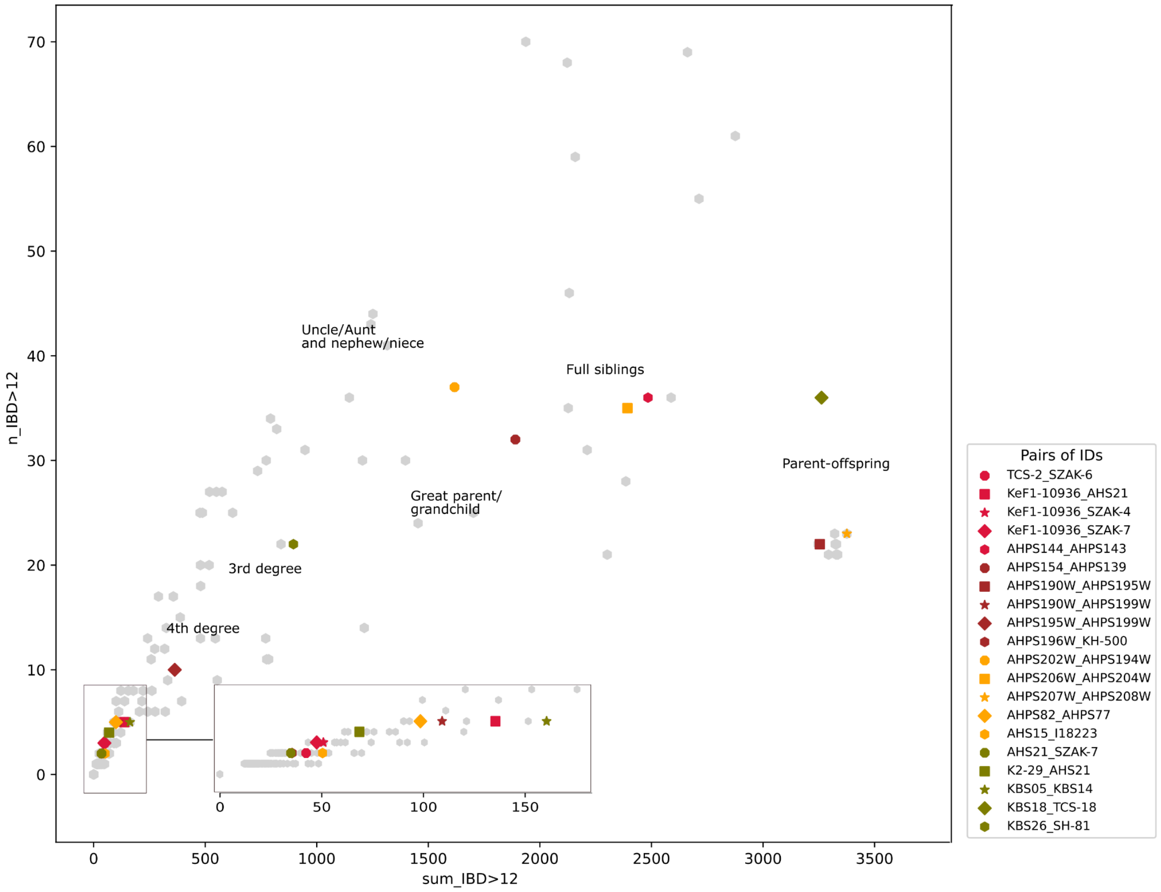


***Figure S26. Scatterplot of IBD (identity-by-descent) segment sharing connections.***
*Relatives detected by IBD within the new dataset, and connections between previously published 5th-12th century individuals (see Main text) and the new samples are highlighted. Laboratory codes and individual codes are matched in Table S1. Kinship types are indicated based on simulations presented in Ringbauer et al.[20].*
